# Climate Change Factors Interactively Shift Peatland Functional Microbial Composition in a Whole-Ecosystem Warming Experiment

**DOI:** 10.1101/2023.03.06.531192

**Authors:** Christopher L. Kilner, Alyssa A. Carrell, Daniel J. Wieczynski, Samantha Votzke, Katrina DeWitt, Andrea Yammine, Jonathan Shaw, Dale A. Pelletier, David J. Weston, Jean P. Gibert

## Abstract

Microbes affect the global carbon cycle that influences climate change and are in turn influenced by environmental change. Here, we use data from a long-term whole-ecosystem warming experiment at a boreal peatland to answer how temperature and CO_2_ jointly influence communities of abundant, diverse, yet poorly understood, non-fungi microbial Eukaryotes (protists). These microbes influence ecosystem function directly through photosynthesis and respiration, and indirectly, through predation on decomposers (bacteria, fungi). Using a combination of high-throughput fluid imaging and 18S amplicon sequencing, we report large climate-induced, community-wide shifts in the community functional composition of these microbes (size, shape, metabolism) that could alter overall function in peatlands. Importantly, we demonstrate a taxonomic convergence but a functional divergence in response to warming and elevated CO_2_ with most environmental responses being contingent on organismal size: warming effects on functional composition are reversed by elevated CO_2_ and amplified in larger microbes but not smaller ones. These findings show how the interactive effects of warming and rising CO_2_ could alter the structure and function of peatland microbial food webs — a fragile ecosystem that stores 25% of terrestrial carbon and is increasingly threatened by human exploitation.

## Introduction

Greenhouse gas emissions set the pace of climate change (1) and most ecosystems on Earth influence this process by serving as carbon (C) sources or sinks (2). Overall, terrestrial ecosystems mitigate*≈* 30 % of anthropogenic CO_2_ emissions (3, 4). The predominant terrestrial carbon sink — peatlands — store 100% more carbon than all forests while covering less than 3% of the globe (5–8). *Sphagnum* mosses dominate peatlands (9) and are responsible for most C sequestration in the form of recalcitrant peat (10). They harbor diverse communities of single-celled (*i*.*e*., bacteria, archaea, protists) and multi-celled (*i*.*e*., fungi, metazoa) organisms (11) that influence C cycling directly (12, 13), through respiration of *≈* 100 Pg · *C* · yr ^*−*1^ in soils alone (14, 15). These organisms indirectly influence the C cycle by supplying nitrogen needed for *Sphagnum* moss growth (16–18), converting methane into the CO_2_ necessary for photosynthesis (19–21), providing stress tolerance to the host mosses (22, 23), and assisting in hostpathogen defense (24).

How ecological interactions between members of the moss-associated community influence their effects on peatland C cycling (25–27), is poorly understood. Predation by metazoans, in particular, plays a prominent role in peatland C cycling (28), and is likely to also impact microbial communities (29, 30). Unicellular Eukaryotes collectively known as protists store two times more C globally than all animals combined (31, 32), contribute to C cycling through mixotrophic metabolism (33), and also serve as one of the principal biological controls on microbial respiration and photosynthesis through predation and competition (30, 34, 35). Protists also directly influence plant growth and health through their effects on the rhizosphere (34, 36, 37) and phyllosphere (38, 39) — thus indirectly influencing plant growth through their effects on beneficial microorganisms (29, 40). In turn, climate effects can induce changes in moss microhabitat that affect predation, resource acquisition, and life cycles of associated protists(33).

But, despite important advances (50–53), how the structure and function of protist communities may respond to climate change in peatlands remains poorly understood. Simple protist functional traits — *i*.*e*., characteristics controlling how organisms respond to the environment or affect ecosystem properties (54) — including protist cell size, cell shape, and cellular contents (Table 1), can be used to predict warming effects on protist communities and ecosystem function (43). This is due to underlying links amongst these traits, metabolism, respiration rates, and energy acquisition mode (43, 44). By quantifying changes in functional traits of these protist communities — in addition to changes in abundances, biomass, and community composition — we can infer changes to their ecological function in these peatland ecosystems. For example, metabolic demand and total respiration rates increase with increasing temperature, favoring smaller organisms (47, 55) with a shape (long and slim) that maximizes energy intake and minimizes energetic costs (42, 56). Increased metabolic demands can impact resource acquisition strategies (e.g., increased heterotrophy over autotrophy) (57–60), resulting in measurable changes in cellular optical properties such as red/green color ratio or the heterogeneity of gray-scale values (*e*.*g*. sigma intensity), which have been linked to cellular contents and cell viability (61, 62). Consequently, changes in functional composition could alter net C fluxes and other elements within peatlands (63–66).

**Table 1.** Functional Justification of Selected Traits Related to Metabolism.

| Trait | Functional Linkage | References |
| --- | --- | --- |
| Cell Size | Cell size determines the metabolic costs of single-celled organisms, and is well-known to decline with temperature through the temperature-size rule. | Brown et al. (41), Atkinson (42), Wieczynski et al. (43) |
| Cell Shape | Cell shape captures information on the surface area to volume ratio of unicellular organisms (i.e., rounder cells have a lower surface/volume ratio than oblong cells), thereby mediating diffusion of material across membranes and establishing metabolic costs of this transport. | Wieczynski et al. (43), Gibert et al. (44), Glazier (45), Hirst et al. (46) |
| Cell Contents | Measured as the optical heterogeneity of a cell, our measure of cellular contents (vacuoles, cytoskeleton, organelles), captures information on cellular storage and metabolism, e.g. *the metabolic machinery of the cell*. | Wieczynski et al. (43) |
| Resource Acquisition Mode | The ratio of the red-to-green reflectance signal of a cell — where higher red-der cells are more likely heterotrophic while greener cells are more likely autotrophic — indicate the resource acquisition mode (heterotrophy, autotrophy, mixotrophy) of the cell, which in turn influences function and C cycling. | Atkinson et al. (47), Gibert et al. (48), Nakano et al. (49) |

Here, we address how peatland protist communities — operationally defined as the community of all non-fungal Eukaryotes — change in novel climates. We quantify shifts in abundance, biomass, taxonomic composition, and functional trait composition in a peatland long-term whole-ecosystem experiment with temperature and CO_2_ manipulations of living moss and superficial peat at the SPRUCE (*Spruce and Peatland Responses Under Changing Environments*) long-term (since 2014) whole-ecosystem warming experimental site (67) (Fig. 1A). We quantified protist abundance, biomass, and community functional traits through fluid imaging and used 18S rRNA gene sequencing to infer mechanisms of change in observed protist community functional trait composition (e.g., through taxonomic shifts or plasticity). We test the hypotheses that increasing temperatures lead to: (1) a decrease in protists’ body size — i.e., the temperature-size rule (42, 47), (2) an increase in surface-to-volume ratio (aspect ratio), as protists optimize shape to increase energy uptake (surface) and reduce metabolic costs (volume) (68), (3) an increase in cellular optical heterogeneity (sigma intensity or cellular “contents”), as temperature-induced increased metabolic rates requires more cellular machinery (69), and (4) increased heterotrophy (increased red/green ratios) to keep up with metabolic demands. Additionally, we expect elevated CO_2_ to result in an increase in total community biomass (70, 71) as more CO_2_ fuels photosynthesis at the base of the food web.

**Fig. 1.**
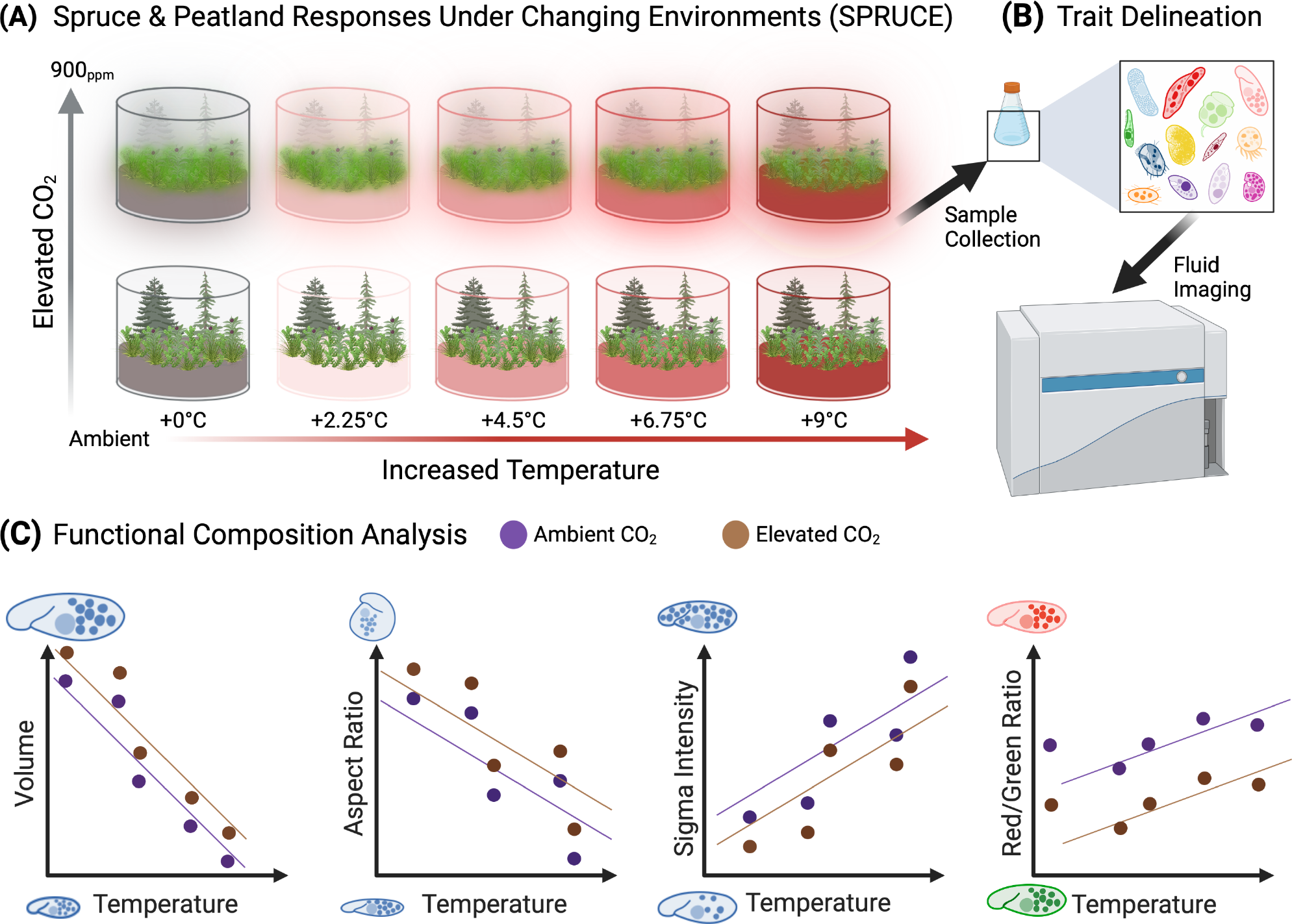
(A) Samples were collected in a 2-factorial mesocosm design in boreal peatlands with variation in temperature (+0, +2.25, +4.5, +6.75, +9°C) and CO_2_ (450 PPM, 900 PPM) concentrations. Protist samples were collected from the upper peat layer within each treatment and processed *ex situ*. (B) After equilibration (24 hours in climate-controlled growth chambers, Carolina protist liquid media, 22°C and 65% humidity), samples were analyzed via fluid imaging to quantify abundances across treatments and physical and optical functional traits. (C) Functional composition data hypothesized on first-principles of metabolic and ecological theory. We anticipate decreases in size (e.g., volume), shape (e.g., aspect ratio), and cellular contents (e.g., sigma intensity) under warming, with an increase in red/green ratio (e.g., photosynthesis). Under elevated CO_2_ we expect larger cell volumes, more roundedness, increased cellular activity, and lower red/green ratios.

This CO_2_ driven increase in energy availability at the base of the microbial food web should also result in (Fig. 1C): an increase in protist volume in response to more energy availability, 2) an increase in aspect-ratio as increased available energy releases cells from shape metabolic constraints, an increase in sigma intensity, due to increased production of cellular machinery to process available energy, and an increase in autotrophy (measured as decreases in the red/green ratio of cells) relative to heterotrophy (47–49). We show that warming and CO_2_ have significant, negative interactive effects, on the taxonomic and functional composition of peatland protist communities that is in turn dependent on organismal size and could result in important consequences for C cycling in peatlands.

## Materials and Methods

### A. Study Site and Field Sampling

The study site is within the S1-Bog of the Marcell Experimental Forest (47° 30.4760^*′*^ N, 93° 27.1620^*′*^ W; 418 m above mean sea level), MN, USA. It sits on the southern edge of the temperate boreal forest range in North America, with a sub-humid continental climate. The S1-Bog — above a water table with marginal groundwater influence — consists of an ombrotrophic peatland dominated by a forest canopy of black spruce (*Picea mariana*) and tamarack (*Larix laricinia*). Understory plants are mixture of ericaceous scrubs, herbaceous sedges and grasses, and a bryophyte layer dominated by *Sphagnum* spp. mosses. Mosses are distributed in hollows (*S. angustifolium & S. fallax*) and in drier hummocks (*S. divinum* previously called *S. magellanicum* and *S. fuscum*). Other bryophytes include feather mosses (*Pleurozium* spp.) and haircap mosses (*Polytrichum* spp.). For further description of the site and below-ground peat profile, see Hanson et al. (2016) and Tfaily et al. (2014).

The *P. mariana*–*Sphagnum* spp. raised bog experimental ecosystem consists of open-top chambers 12m in diameter in a factorial design of ambient 450 ppm and elevated 900 ppm CO_2_ treatments across ambient (+0, +2.25, +4.5, +6.75, +9°C) soil (initiated June 2014) and air (initiated June 2015) warming treatments (Fig. 1; (72)). After 5 years of treatment — in September 2019 — we collected superficial samples (1-5 cm) from SPRUCE containing living plant tissue and superficial peat from two sub-plot locations within each enclosure. We limited sampling efforts to a minimally sufficient amount of material to avoid disturbing the overall peatland community and interfering with long-term experiments at SPRUCE. We placed the samples in individual sterile bags and over-night shipped them to Durham, NC for imaging of protist communities while additional samples were flash-frozen and over-night shipped to Oak Ridge, TN for DNA extraction and 18S amplicon sequencing. Upon reception of samples for imaging, we removed the material from sterile bags, placed them in cloth sieves within glass funnels, and washed them thoroughly with Carolina protist media (Liquid Protozoan Nutrient Concentrate; 13-2350). We collected the discharge from the funnels in 250 ml glass jars and kept the cultures at 22°C and 65% humidity for equilibration for 24 hours before fluid imaging. All samples were processed and imaged in a simultaneous, treatment-blind fashion, and within 24-36 hours of collection in the field. While samples could have been fixed *in situ* with lugol or formaldehyde, these fixative methods often alter the size and shape of cells and lead to an under-representation of larger size classes (74, 75), and would bias our quantification of functional trait shifts across size classes with temperature and CO_2_.

### B. Density and Biomass

Protists were operationally defined in this study as all non-fungal unicellular Eukaryotes. Their densities were estimated as simple counts 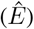 of the number of imaged particles per volume of fluid analyzed through fluid imaging with a FlowCAM (Yokagawa Fluid Imaging, USA) at 10*×* magnification of 6 mL water sub-samples of processed SPRUCE samples. The FlowCam screens roughly 75% of the sampled volume (called, the “efficiency” of the machine), which was used to efficiency-correct biomass and abundance estimates from total protist volume and densities (see supplementary materials; Fig. S2).

### C. Trait Measurements

We quantified functional traits (Table S1) by manually curating the ensuing cell images in Visual Spreadsheet software program (Yokagawa Fluid Imaging, USA), discarding all non-protist images, cysts, and other organic materials and debris, and retaining only high-quality protist images for trait quantification. All protist images (n = 211,039) were used to quantify densities. To test our hypotheses about size, shape, cellular contents, and energy acquisition mode, we used four traits (volume, aspect ratio, sigma intensity, red/green ratio) representing these functional trait categories from a larger set of 37 physical and optical traits; these representative traits were chosen due to their linkages to the functions of interest (Table 1) and by eliminating highly correlated and co-linear traits (see supplementary materials; Table S1, Fig. S1A). Wieczynski et al. (2021) found these four traits to influence temperature performance curves (TPCs) of protists and their metabolic functioning (Table 1).

### D. Regression Analyses

We tested the effects of temperature and CO_2_ levels on the abundance and biomass of the protist communities with generalized additive models (GAMs) and used generalized linear models (GLMs) to test for effects on the traits. We pooled the data from each enclosure subplot, as the level of inference is that of the enclosure (Fig. 1). To test hypotheses 1-4, we first clustered individual cellular observations into size classes based upon the natural log of protists’ geodesic length (e.g., longest axis) using Gaussian finite mixture modelling with the R package *mclust* (76). We chose length as it was less correlated with the traits of interest and would reduce co-linearity in our models between the size class cluster and the functional traits (Fig. S1). We selected an optimal set of five size classes with a Gaussian mixture model and Bayesian Information Criterion (BIC) (Fig. S3). We then ran GLMs for each functional trait against size class (factor), temperature (continuous), and carbon dioxide level (factor). Associated Pearson correlation coefficients and p-values were obtained via the *ggpubr* R package (77) and are reported for each size class (Fig. S4). Additionally, we examined changes across environments in the estimated abundance and biomass (grams) of the community per size class with GAMs (k = 5) using the R package *mgcv* (78), reporting associated Pearson correlations and p-values for each size class (Fig. S5).

### E. Structural Equation Modeling (SEM) and Functional Diversity

To quantify possible direct and indirect effects of Temperature and CO_2_ on protist communities, we used Structural Equation Modeling (SEM) with the R package *lavaan* (79), a robust Maximum Likelihood (MLM) estimator, and the ‘csolnp’ optimizer for enhanced convergence properties. The first SEM considered the effects of the environment (Temperature, CO_2_, and their interaction) on compositional (Observed Richness, Shannon Diversity, and Biomass) and functional trait diversity (Aspect Ratio, Red/Green Ratio, Sigma Intensity, and Volume), and also accounted for possible effects of community composition and structure on functional trait diversity. A second SEM included latent variables representing compositional (Observed Richness, Shannon Diversity) and functional trait diversity (Aspect Ratio, Red/Green Ratio, Sigma Intensity, and Volume). A bootstrap procedure with 1000 replications was used to estimate robust standard errors and p-values for all model effects. We assessed model fit through conventional indices for goodness-of-fit (Comparative Fit Index (CFI), Tucker-Lewis Index (TLI), Standardized Root Mean Square Residual (SRMR), and Root Mean Square Error of Approximation (RMSEA)). To quantify the functional diversity of our protist communities, we used the *FD* and *fundiversity* packages in R (80, 81), leveraging the mean trait values for each size class within treatments.

### F. Amplicon Sequencing

To understand the possible mechanisms behind observed changes in abundance, biomass and functional traits, we extracted genomic DNA from our samples with the DNeasy 96 Plant Kit (Qiagen) and amplified the V4 region of the 18S rRNA gene following (82). Samples were multiplexed, pooled and sequenced on an Illumina MiSeq instrument with paired end kit (2 x 250 bp). We processed protist sequences with the QIIME 2 v 2021.2 platform. Paired sequences were first demultiplexed with the plugin demux; then, we quality filtered (denoised, dereplicated, chimera filtered and pair-end merged) and processed in Sequence Variants (SVs) with the dada2 plugin (83). Taxonomy followed the Pr2 (18S) database (84) and sequences assigned as “bacteria” or Embryophyceae were removed. We used sequence variants to calculate alpha diversity as counts and the Shannon diversity (calculated beta diversity) and compositional dissimilarity, or Bray-Curtis distance, with the *phyloseq* package (85).

## Results

Experimental manipulations of temperature and CO_2_ levels induced an 80-fold change (+/-4.45%) in total observed protist densities (F = 2.2554; p = 0.02244, Fig. S5A) and biomass (F = 2.2504; p = 0.02273, Fig. S5B) across treatments (Fig. 2A), as well as a shift in the relative dominance of size classes (Fig. 2B). Across all body size classes, estimated protist density and biomass increased with temperature under ambient CO_2_ (Fig. 2A). This trend reversed under elevated CO_2_, with major declines in biomass and density across size classes (Fig. 2A). The relative density within size classes remained generally consistent with increasing temperature under ambient CO_2_ conditions; however, under elevated CO_2_ conditions, the relative density of larger protists increased at the expense of the two smallest size classes (Fig. 2B).

**Fig. 2.**
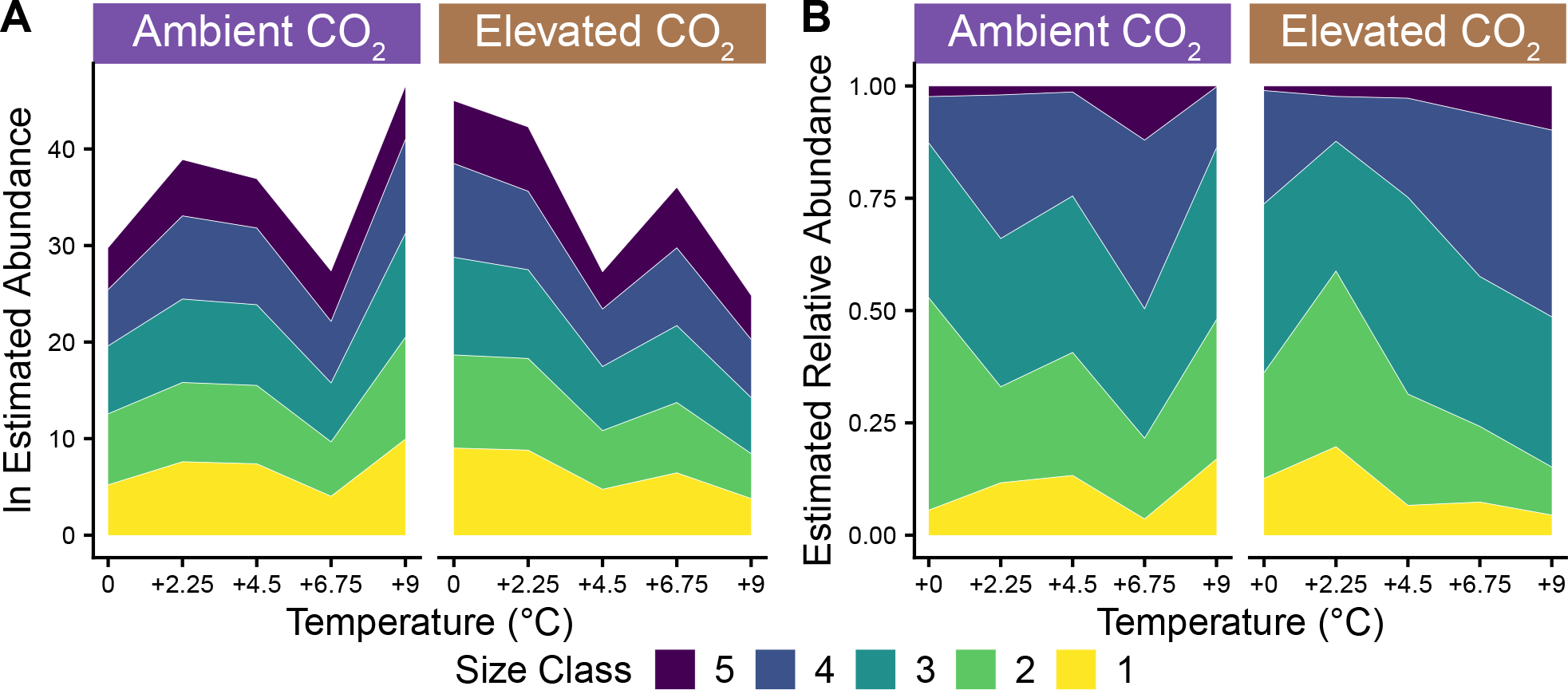
Estimated protist abundances derived from FlowCam densities. (A) Estimated total abundance data on the natural log scale for size classes across temperatures under ambient and elevated CO_2_ treatments. Error rates for all size classes are +/1 4.45%. Data indicates that abundances increase across size classes with temperature under ambient CO_2_ levels but decreases under elevated CO2 levels. (B) Relative abundance of size classes by CO_2_ treatment calculated from total estimated abundance (A) across temperatures. Results show that different size classes respond differentially to environmental treatments. Size Class 1: 12.43 *μ*m - 17.07 *μ*m; Size Class 2: 17.07 *μ*m - 20.68 *μ*m; Size Class 3: 20.68 *μ*m - 28.24 *μ*m; Size Class 4: 28.24 *μ*m - 59.74*μ*m; Size Class 5: 59.74 *μ*m - 526.43 *μ*m.

A three-way interaction between temperature, CO_2_ concentration, and size class drove significant changes in the functional trait composition of protist communities (Fig. 3): size (F = 4,007; *p ≪* 2.2*x*10^*−*16^); shape (F = 13,204; *p ≪* 2.2 *×* 10^*−*16^); cellular contents (F = 789.05; p *≪* 2.2*x*10^*−*16^); and energy acquisition (F = 657.15; p *≪* 2.2*x*10^*−*16^). The magnitude of the change was driven by the CO_2_ treatment, with elevated CO_2_ rates of change outpacing that of ambient CO_2_ by up to a factor of 25 (Fig. 3). Under ambient CO_2_, protists were smaller (Figs. 3A, S6), less round (Figs. 3B, S7), showed more cellular contents (Figs. 3C, S8), and became redder (more heterotrophic) at lower levels of warming but greener (more autotrophic) at higher levels of warming (Fig. 3D, Fig. S9). Conversely, protists under elevated CO_2_ were larger (Fig. 3A, S6), rounder (Fig. 3B, S7), had fewer cellular contents (Fig. 3C, S8), and got redder (Fig. 3D, S9). Thus, with the exception of the two smallest size classes for red/green ratio, protists displayed inverse patterns for all traits in response to increasing temperature under ambient and elevated CO_2_ levels.

**Fig. 3.**
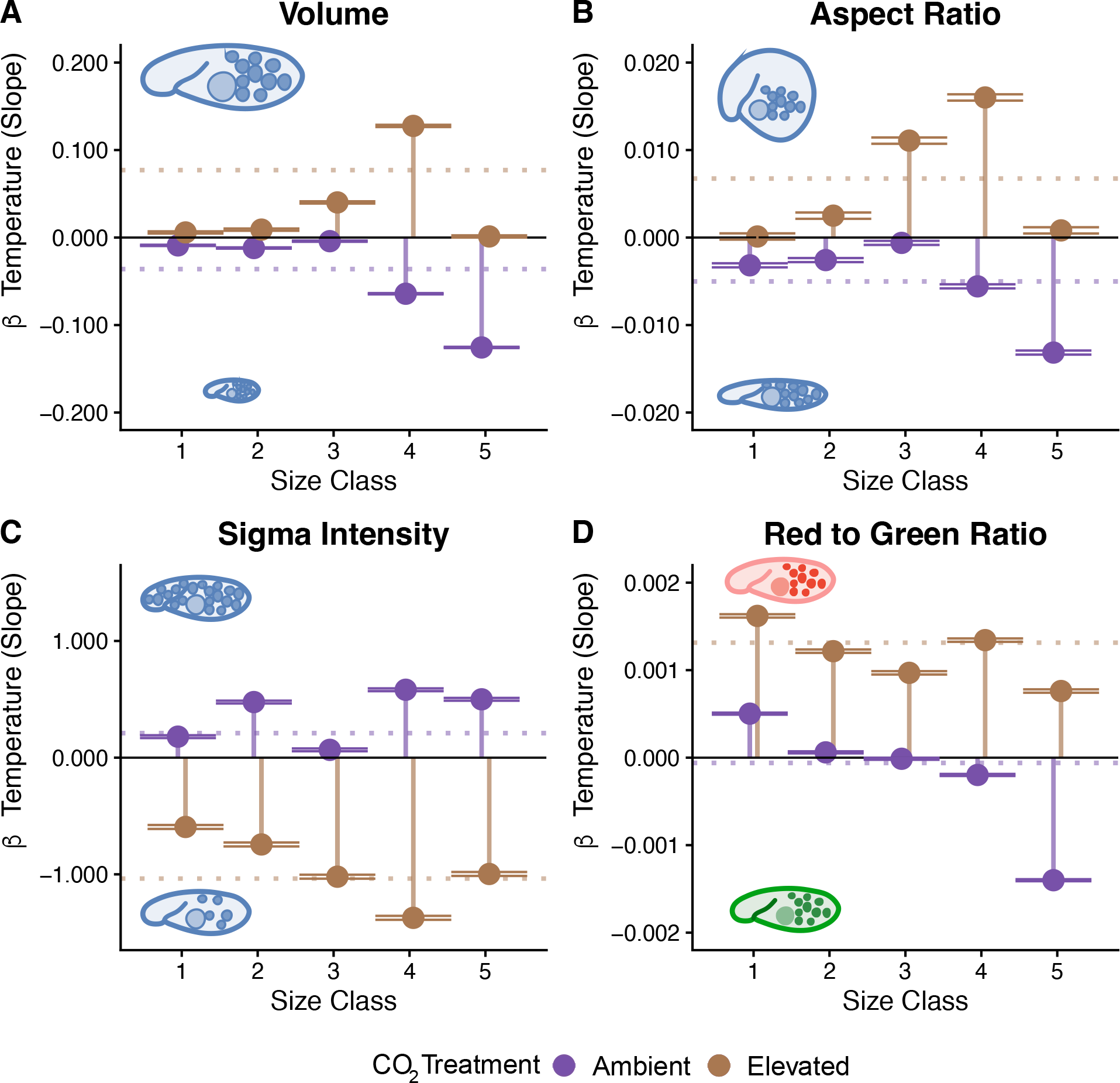
Rate of change across temperatures (*β* or slope) for functional traits by protist size class and CO_2_ treatments (lollipops). Slopes were calculated from GLMs. Mean slope across size classes displayed as dashed line. Positive slope values indicate: (A, Volume) increasing size; (B, Aspect Ratio) higher symmetry and roundedness (shape); (C, Sigma Intensity) higher optical heterogeneity; and (D, Red/Green ratio) increasing redness and/or decreasing greenness.

The observed changes in functional traits were protist-size dependent, as larger size classes responded more strongly to temperature than smaller ones (Fig. S10A, (0.014 *<* p *<* 0.066), c.f. (55)). While the rates of change in all traits displayed non-linear size-dependency, we found weak support for size allometries in shape, cellular contents, and energy acquisition (Fig. S10B-D).

Observed changes in community functional traits could be the result of plastic change across all or most protists in the community, or, instead, result from taxonomic compositional shifts. Amplicon sequencing results show that important taxonomic shifts (Fig. 4A) likely underlie the observed changes in functional traits (Fig. 3), not species-specific plastic change. Moreover, increasing temperature leads to a decrease in the diversity of microbial communities (Fig. 4B,C) — consistent with multiple studies across biomes that find climate change decreases diversity of species across scales (86–88). Importantly, temperature interacts with CO_2_ such that the effects of CO_2_ treatment decrease at warmer temperatures, leading to a near convergence in the taxonomic compositional similarity between ambient and elevated CO_2_ (Fig. 4D), despite a dramatic divergence in functional traits (Fig. 3).

**Fig. 4.**
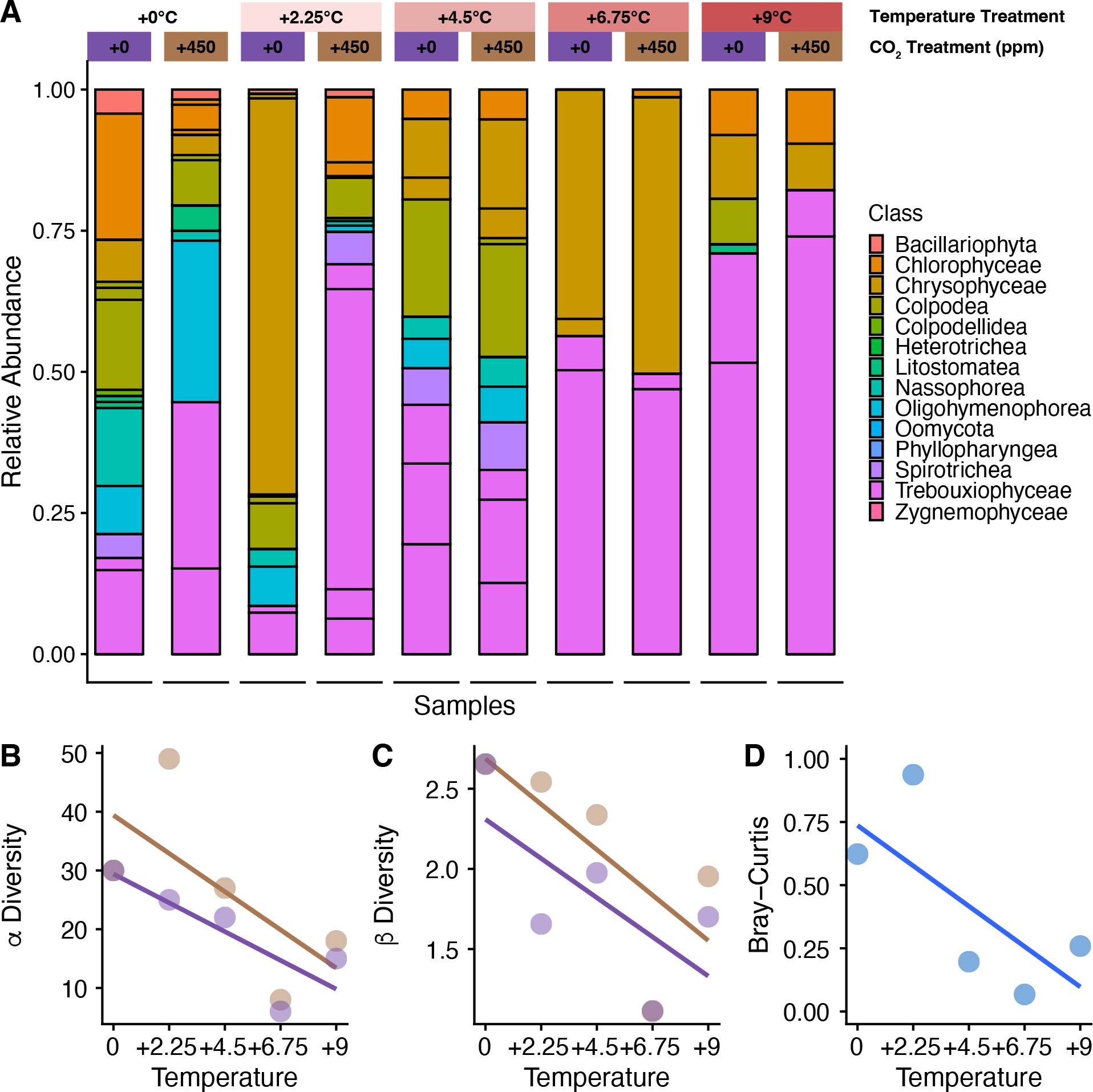
Change in (A) protist Order relative composition (colored by Class) and (B, C) diversity of protist community across temperatures and CO_2_ treatments. (A) Protists’ Order composition (colored by Class) shifts both with temperature increase and within CO_2_ treatment, with a decrease in (B) total species composition and (C) species diversity at higher temperatures. (D) Species composition becomes more homogeneous — lower Bray-Curtis distance — between CO_2_ treatments at higher temperatures, indicating an interactive effect (pearson’s correlation = −0.71).

Our SEM model adequately described the data (CFI = 0.96, TLI = −0.771, SRMR = 0.025), and supported the existence of strong direct interactive effects of temperature and CO_2_ on community structure and composition and community functional traits (Fig. 5A, Table S3). The model revealed stronger pressure on individual traits and community diversity from CO_2_ than temperature — though their interactive effects weighed most heavily on both traits and community structure metrics (Fig. 5A). Moreover, we uncovered the existence of indirect effects of temperature, CO_2_, and their interaction on community functional traits, through direct effects on species richness, which in turn influences cell size, shape, contents and energy acquisition mode (Figs. 5, S12).

**Fig. 5.**
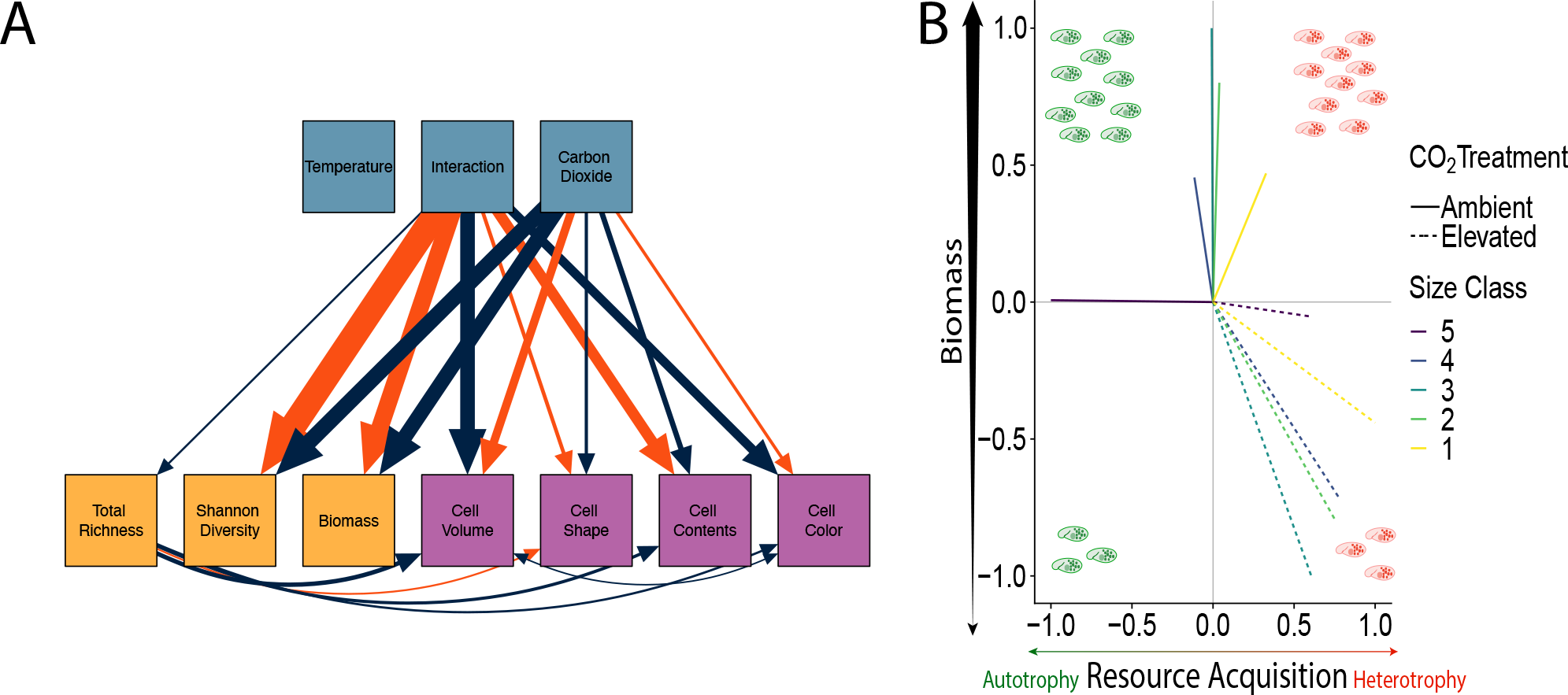
(A) Structural Equation Model (SEM) illustrating the effects of environmental variables and their interaction (light blue) on protist community structure (light orange) and functional traits (purple). Solid lines — shown only the top 30% stronger effects — represent significant effects, with positive effects (blue) and negative effects (red). The thickness of the lines corresponds to the magnitude of the standardized estimates (Table S3). Traits are as follows: Shape = Aspect Ratio, Color = Red to Green Ratio, Cell Contents = Sigma Intensity. (B) Combined normalized change in biomass and resource acquisition across quadrant (I-IV) outcomes. Under elevated CO_2_, the increase in respiration is offset by a loss in biomass of these organisms (IV). Given normalization of difference, slopes *≤* −1 indicate a net reduction in heterotrophy. Under ambient CO_2_ conditions, an increase in net photosynthesis occurs for the two largest size classes (II), as both biomass and green reflectance increase. The net carbon dynamics in these plots may be offset by the normalized relative increase in biomass of the more heterotrophic smaller size classes (I).

These indirect effects often had a reverse sign, suggesting small possible offsets through changes in total protist richness. For example, the interaction between temperature and CO_2_ reduces protist size directly, but indirectly, it also can increase protist size through reducing richness, which itself negatively influences cell size (Fig. 5A). These results held while accounting for variations in the model structure of the SEM (i.e., including latent variables, Table S2, Fig. S12).

Lastly, functional diversity analysis supported the conclusion that temperature and CO_2_ have complex joint effects on the structure of protist communities that diverge from the direct effects of each factor independently (Fig. S13). Surprisingly, we found no relationship between Functional Richness or Evenness and Temperature across CO_2_ treatments (Fig S13A, B). We did, however, find a steep decline in functional dispersion with rising temperature under elevated CO_2_, whereas the ambient CO_2_ scenario showed a more moderate response (Figure. S13C).

## Discussion

Understanding how moss-associated microbial communities change with temperature and CO_2_ concentrations is vital for predicting how peatland ecosystems will respond to climate change (7, 89, 90) and alter the global C cycle (91–93). Here we find strong evidence of changes in the abundance, biomass, functional traits, and composition of important but poorly understood members of peatland microbial communities. These changes are driven by an interaction between warming and CO_2_ concentration (Figs. 2-5). We show that peatland protist communities are sensitive to climate change, and under ambient CO_2_ match our expectations based upon metabolic theory — getting smaller, less round, more active, and more heterotrophic. However, these trends are reversed by elevated CO_2_ concentrations (Fig. 3), and the magnitude and direction of these changes are driven by non-linear size-dependence (Fig. S10). While protist communities converge in taxonomic composition between ambient and elevated CO_2_ levels as temperature rises, they diverge in functional trait composition (Figs. 3, 4, S13, S14), suggesting important but yet unknown effects on peatland function with environmental change.

Protist reductions in volume (Figs. 3A, S6) and roundedness (Fig. 3B, Fig. S7) observed under ambient CO_2_ and elevated temperatures were predicted from metabolic theory, suggesting that we can anticipate complex temperature responses in natural and intricate microbial communities based on first principles. For example, temperature selects for smaller organisms and/or organisms with larger surface-to-volume ratios that can meet higher metabolic demands while conserving resources as metabolism costs increase at higher temperatures (94). This negative relationship between size and temperature — the “temperature-size rule” (42, 47) — is well documented for both single-celled (47) and multi-celled organisms (95–97), but was so far thought to be a plastic response.

We show that the temperature-size rule can result from community composition shifts that select for smaller organisms (Figs. 3-5). Similarly, shape shifts (e.g., a decrease in roundedness with temperature) likely results from selection to conserve energy by simultaneously reducing volume (which increases metabolic costs), and increasing surface (which increases resource acquisition) (98). Changes in protist shape and size have multiple ecological consequences, including population dynamics and ecological interactions within communities (30, 43, 44), as well as total respiration, which decreases with volume (99). Shifts in size and shape can occur quite rapidly and can be as dramatic as 50% or more (48, 100). Likely, species that cannot rapidly respond to temperatures are replaced by those that can, leading to high species turnover (Fig. 4A).

The increase in the red/green ratio of 70% of size classes in our study (Figs. 5B, S9) may be alarming. A decrease in the red/green ratio is associated with decreased autotrophy and therefore carbon uptake (101). However, when we normalize the *β* coefficents of our models, the overall increase in heterotrophy is outweighed by a decrease in biomass of heterotrophic organisms and an increase in biomass of photosynthetic organisms obtained through our 18S data (Fig. S14), likely indicating a net reduction in heterotrophy (Figs. 5B,S9, S12). The results of this increase in the green biomass may be linked to higher abundance (Fig. 4A) and shifts in the functional metabolic traits (Figs. 5A, S14, Table S4) of photosynthetic clades such as Trebouxiophyceae; this increase in phototrophic protists with warming — modulated by CO_2_ — parallel to shifts in their traits suggests a shift in metabolic activity of the community, a fundamental result in light of the importance of phototrophic eukaryotes for both the peatland C cycle (102, 103) and the overall global carbon cycle (104). Coupled with decreases in the abundance of consumer microbes like Oligohymenophorans and Oomycota (Fig. 4A), our findings point to potential shifts in the functional metabolome and carbon cycling of peatlands under climate change (Fig. 5B, Fig. S14, Table S4).

Counter to our hypothesis, we report surprising reversals in functional trait change with temperature between CO_2_ treatments (Figs. 3, 4, S6-S9) suggesting that joint temperature and CO_2_ impacts in microbial food webs may be more complex than previously thought (Figs. 5, S12). Interactive effects of CO_2_ and temperature have been reported in aquatic phytoplankton systems (70, 71, 105). However, these systems only displayed shifts in trait change magnitudes, not complete trend reversals, as reported here. Our results thus emphasize the importance of addressing temperature impacts on microbial organisms in the light of simultaneous CO_2_ increases. These trend reversals may partially be due to CO_2_-induced pH changes of the peat and/or the stoichiometry of diffusion under higher temperatures and elevated CO_2_. Experimental work on ocean phytoplankton showed that rising temperatures and elevated CO_2_ can induce a more rapid decrease in community biomass and chlorophyll-a concentration (106), and that elevated CO_2_ may lessen the demand for a higher surface-to-volume ratio as diffusion becomes easier across the cellular membrane at higher concentrations of dissolved CO_2_. These changes may also be due to the increased efficiency of heterotrophic resource acquisition compared to photosynthesis (107). The shift towards heterotrophic resource acquisition (Fig. 3D) under elevated CO_2_ conditions may lessen the need for energy conservation.

These reported trend reversals under elevated CO_2_ may also occur indirectly, through changes in nutrient cycling and mineralization (108) — especially of N (109). Elevated CO_2_ often increases the exudation of soluble organic compounds by *Sphagnum* mosses (110) — especially sugars — that contribute to the structure of microbial communities, and can increase the abundance of bacteria and heterotrophic protists (111). With higher abundances of bacteria, protists may obtain more energy through predation. Petro et al. (2023) recently found similar interactive effects of warming and elevated CO_2_ on N. Elevated temperature and ambient CO_2_ conditions produced accumulation of nitrogen and a reduction in N_2_ fixation, which was reversed under elevated CO_2_ conditions. This in turn altered N immobilization by micro-organisms, likely leading to a change in microbial food web dynamics. Elevated CO_2_ may also indirectly increase nitrogen availability through changes in *Sphagnum* exudation of C and N organic compounds (26, 113) — indirectly influencing N-releasing enzymes (114, 115) — or through symbiosis with mycorrhizal fungi (116, 117).

We found that overall, the trend reversal reported here (biomass, abundance, traits) was size-dependent (Fig. S10). We find strong support for a super-linear scaling relationship between body size and volume that indicates protist populations might undergo density-dependent (population) and density-independent (individual phenotypic) responses to climate change (43, 48, 100, 118, 119). Our results indicate a significant relationship between proportional changes in traits and size. Larger protists’ size (Fig. S10A) and shape (Fig. S10B) change more than smaller protists per unit volume (Figs. S10A, S10B). Cellular contents follow a sub-linear size-dependency (Fig. S10C). Because smaller protist phenotypes shift much less and these smaller individuals dominate the relative abundance of the system (Figs. 2B, S3B), there may be embedded resistance to phenotypic change in these communities overall. This trend portends potential eco-evolutionary responses of protists under environmental stressors, as phenotypic traits and plasticity may drive the success and survival of individuals responding to changing metabolic demands (120, 121).

Interestingly, our SEM model (Fig. 5A, Tables S2, S3) uncovers additional mechanisms through which these trend reversals are possible. For example, all SEM models found indirect pathways from community structure and composition to functional traits that reversed the direct effects of environmental conditions on functional trait changes (Figs. 5A, S11, S12). While these indirect effects are weaker in magnitude than the direct effects, they also somewhat offset the direct effects through trend reversals (Fig. 5A), perhaps shielding the community from even larger community functional trait shifts. Understanding these complex responses is pivotal for predicting how ecosystem services, such as nutrient cycling and primary productivity, might be reshaped in a future, warmer and CO_2_-enriched world (122–124).

### G. Caveats

Due to the limited temporal resolution of our sampling, it is possible that our results are only indicative of large shifts in the seasonal or transient dynamics of these moss-associated protist communities. However, given the magnitude of the observed changes, we suspect — but do not prove — that these reported shifts owe to broader non-transient community responses to temperature and CO_2_ levels. It is also possible that our sample processing may have inserted bias in our protist counts across temperature treatments. This is because the equilibration period between extraction of the community and fluid imaging — roughly 24 hours at room temperature — could have deferentially affected our samples across treatments. However, we argue that such an effect should have been small, if at all present, as SPRUCE experiences major daily temperature fluctuations in September — when these data were collected — with daily temperatures that can range from below freezing to above 30°C (National Weather Service: https://www.weather.gov). If our samples had changed in composition so dramatically that our data could not be properly interpreted, it would also suggest that SPRUCE protist communities experience extremely large daily changes in composition. Such changes are likely on the longer timescales of months and seasons (125–127), but have, to our knowledge, not been shown to occur in the timescale of days. Last, because our density data was not controlled by total dry moss biomass — known to be positively correlated with microbial abundances (128) — it may show unintended levels of variability. However, we have designed our sampling regime so that living and dead moss biomass was consistent across samples and enclosures, so the amount of variation should be small. Also, the density data recovers patterns of temperature and CO_2_ interactions that are consistent with those of trait and 18S data, both of which are unaffected by the actual amounts of dry moss biomass. Because of this, we acknowledge the possibility that these three sources of error may have affected our results, but remain confident that our inference is robust.

## Conclusions

Our findings suggest important ways in which climate change may influence C cycling in peatland ecosystems by potentially altering important functional linkages within moss-associated microbial food webs (129). Further, while earth-system models may predict range shifts in *Sphagnum* spp. (130), our findings indicate that the combined effects of elevated carbon dioxide and temperature may alter the symbiotic moss-microbiome in unexpected ways — ways that could affect the distribution of *Sphagnum* and peatlands in the future (23, 26, 27, 112). Given the diverse energy acquisition strategies of protists, changes in the balance of autotrophy versus heterotrophy can have cascading effects on the biological carbon cycle (131–134). Due to their predominant distributions in mid- and high-latitudes of the Northern hemisphere, peatlands are especially sensitive to climate change (135–138). While tantalizing, the data-driven hypotheses presented here need to be tested further, but are suggestive of important ways in which environmental change both influences and is influenced by rapid microbial responses in functional trait composition in globally important — and highly threatened — ecosystems such as peatlands.

## AUTHOR CONTRIBUTIONS

CLK and JPG conceptualized the study. AC, DAP and DJW collected SPRUCE samples. SV, AY and JPG prepared the samples and extracted protist data. AAC prepared and provided amplicon sequencing data. CLK performed all analyses with support from DJW, AC, and JPG. CLK wrote the first draft and all authors contributed substantially to subsequent versions.

## ACKNOWLEDGEMENTS

We would like to thank Dawn M. Klingeman from Oak Ridge National Laboratory for running the 18S rRNA samples on the sequencer and two anonymous reviewers for their feedback.

## Bibliography

1. Will Steffen, Johan Rockström, Katherine Richardson, Timothy M Lenton, Carl Folke, Diana Liverman, Colin P Summerhayes, Anthony D Barnosky, Sarah E Cornell, Michel Crucifix, et al. Trajectories of the earth system in the anthropocene. Proceedings of the National Academy of Sciences, 115(33):8252–8259, 2018.

2. Benjamin D Stocker, Raphael Roth, Fortunat Joos, Renato Spahni, Marco Steinacher, Soenke Zaehle, Lex Bouwman, Iain Colin Prentice, et al. Multiple greenhouse-gas feed-backs from the land biosphere under future climate change scenarios. Nature Climate Change, 3(7):666–672, 2013.

3. Josep G Canadell, Corinne Le Quéré, Michael R Raupach, Christopher B Field, Erik T Buitenhuis, Philippe Ciais, Thomas J Conway, Nathan P Gillett, RA Houghton, and Gregg Marland. Contributions to accelerating atmospheric co2 growth from economic activity, carbon intensity, and efficiency of natural sinks. Proceedings of the national academy of sciences, 104(47):18866–18870, 2007.

4. J.G. Canadell, P.M.S. Monteiro, M.H. Costa, L. Cotrim da Cunha, P.M. Cox, A.V. Eliseev, S. Henson, M. Ishii, S. Jaccard, C. Koven, A. Lohila, P.K. Patra, S. Piao, J. Rogelj, S. Syampungani, S. Zaehle, and K. Zickfeld. Global Carbon and other Biogeochemical Cycles and Feedbacks, page 673–816. Cambridge University Press, Cambridge, United Kingdom and New York, NY, USA, 2021. doi: 10.1017/9781009157896.007.

5. Florian Humpenöder, Kristine Karstens, Hermann Lotze-Campen, Jens Leifeld, Lorenzo Menichetti, Alexandra Barthelmes, and Alexander Popp. Peatland protection and restoration are key for climate change mitigation. Environmental Research Letters, 15(10): 104093, 2020.

6. Jiren Xu, Paul J Morris, Junguo Liu, and Joseph Holden. Peatmap: Refining estimates of global peatland distribution based on a meta-analysis. Catena, 160:134–140, 2018.

7. J Loisel, Angela V Gallego-Sala, and Z Yu. Global-scale pattern of peatland sphagnum growth driven by photosynthetically active radiation and growing season length. Biogeosciences, 9(7):2737–2746, 2012.

8. Zicheng Yu, Julie Loisel, Daniel P Brosseau, David W Beilman, and Stephanie J Hunt. Global peatland dynamics since the last glacial maximum. Geophysical research letters, 37(13), 2010.

9. Merritt R Turetsky, Benjamin Bond-Lamberty, E Euskirchen, Julie Talbot, Steve Frolking, A David McGuire, and E-S Tuittila. The resilience and functional role of moss in boreal and arctic ecosystems. New Phytologist, 196(1):49–67, 2012.

10. Aljosja Hooijer, Susan Page, JG Canadell, Marcel Silvius, Jaap Kwadijk, H Wösten, and Jyrki Jauhiainen. Current and future co 2 emissions from drained peatlands in southeast asia. Biogeosciences, 7(5):1505–1514, 2010.

11. Joel E Kostka, David J Weston, Jennifer B Glass, Erik A Lilleskov, A Jonathan Shaw, and Merritt R Turetsky. The sphagnum microbiome: new insights from an ancient plant lineage. New Phytologist, 211(1):57–64, 2016.

12. D Gilbert, Christian Amblard, Gilles Bourdier, and A-J Francez. The microbial loop at the surface of a peatland: structure, function, and impact of nutrient input. Microbial ecology, 35(1):83–93, 1998.

13. Daniel Gilbert and Edward AD Mitchell. Microbial diversity in sphagnum peatlands. Developments in earth surface processes, 9:287–318, 2006.

14. Christos Gougoulias, Joanna M Clark, and Liz J Shaw. The role of soil microbes in the global carbon cycle: tracking the below-ground microbial processing of plant-derived carbon for manipulating carbon dynamics in agricultural systems. Journal of the Science of Food and Agriculture, 94(12):2362–2371, 2014.

15. Susan Trumbore. Carbon respired by terrestrial ecosystems–recent progress and challenges. Global Change Biology, 12(2):141–153, 2006.

16. Andreas Berg, Åsa Danielsson, and Bo H Svensson. Transfer of fixed-n from n2-fixing cyanobacteria associated with the moss sphagnum riparium results in enhanced growth of the moss. Plant and soil, 362(1):271–278, 2013.

17. Melanie A Vile, R Kelman Wieder, Tatjana Živković, Kimberli D Scott, Dale H Vitt, Jeremy A Hartsock, Christine L Iosue, James C Quinn, Meaghan Petix, Hope M Fillingim, et al. N2-fixation by methanotrophs sustains carbon and nitrogen accumulation in pristine peatlands. Biogeochemistry, 121(2):317–328, 2014.

18. Kathrin Rousk, Davey L Jones, and Thomas H DeLuca. Moss-cyanobacteria associations as biogenic sources of nitrogen in boreal forest ecosystems. Frontiers in Microbiology, 4: 150, 2013.

19. Nardy Kip, Wenjing Ouyang, Julia van Winden, Ashna Raghoebarsing, Laura van Niftrik, Arjan Pol, Yao Pan, Levente Bodrossy, Elly G van Donselaar, Gert-Jan Reichart, et al. Detection, isolation, and characterization of acidophilic methanotrophs from sphagnum mosses. Applied and environmental microbiology, 77(16):5643–5654, 2011.

20. Nardy Kip, Christian Fritz, ES Langelaan, Yao Pan, Lev Bodrossy, Veronica Pancotto, MSM Jetten, AJP Smolders, and HJM Op den Camp. Methanotrophic activity and diversity in different sphagnum magellanicum dominated habitats in the southernmost peat bogs of patagonia. Biogeosciences, 9(1):47–55, 2012.

21. Max Reumer, Monika Harnisz, Hyo Jung Lee, Andreas Reim, Oliver Grunert, Anuliina Putkinen, Hannu Fritze, Paul LE Bodelier, and Adrian Ho. Impact of peat mining and restoration on methane turnover potential and methane-cycling microorganisms in a northern bog. Applied and environmental microbiology, 84(3):e02218–17, 2018.

22. AV Shcherbakov, AV Bragina, E Yu Kuzmina, Christian Berg, AN Muntyan, NM Makarova, NV Malfanova, Massimiliano Cardinale, Gabriele Berg, VK Chebotar, et al. Endophytic bacteria of sphagnum mosses as promising objects of agricultural microbiology. Microbiology, 82(3):306–315, 2013.

23. Alyssa A Carrell, Travis J Lawrence, Kristine Grace M Cabugao, Dana L Carper, Dale A Pelletier, Jun Hyung Lee, Sara S Jawdy, Jane Grimwood, Jeremy Schmutz, Paul J Hanson, et al. Habitat-adapted microbial communities mediate sphagnum peatmoss resilience to warming. New Phytologist, 234(6):2111–2125, 2022.

24. Katja Opelt, Vladimir Chobot, Franz Hadacek, Susan Schönmann, Leo Eberl, and Gabriele Berg. Investigations of the structure and function of bacterial communities associated with sphagnum mosses. Environmental microbiology, 9(11):2795–2809, 2007.

25. Verity G Salmon, Deanne J Brice, Scott Bridgham, Joanne Childs, Jake Graham, Natalie A Griffiths, Kirsten Hofmockel, Colleen M Iversen, Terri M Jicha, Randy K Kolka, et al. Nitrogen and phosphorus cycling in an ombrotrophic peatland: a benchmark for assessing change. Plant and Soil, 466(1):649–674, 2021.

26. Alyssa A Carrell, Max Kolton, Jennifer B Glass, Dale A Pelletier, Melissa J Warren, Joel E Kostka, Colleen M Iversen, Paul J Hanson, and David J Weston. Experimental warming alters the community composition, diversity, and n2 fixation activity of peat moss (sphagnum fallax) microbiomes. Global Change Biology, 25(9):2993–3004, 2019.

27. Alyssa A Carrell, Dušan Veličković, Travis J Lawrence, Benjamin P Bowen, Katherine B Louie, Dana L Carper, Rosalie K Chu, Hugh D Mitchell, Galya Orr, Lye Meng Markillie, et al. Novel metabolic interactions and environmental conditions mediate the boreal peatmoss-cyanobacteria mutualism. The ISME journal, 16(4):1074–1085, 2022.

28. Kevin H Wyatt, Kevin S McCann, Allison R Rober, and Merritt R Turetsky. Trophic interactions regulate peatland carbon cycling. Ecology Letters, 24(4):781–790, 2021.

29. Stefan Geisen, Enrique Lara, Edward AD Mitchell, Eckhard Völcker, and Valentyna Krashevska. Soil protist life matters! Soil Organisms, 92(3):189–196, 2020.

30. Jennifer D Rocca, Andrea Yammine, Marie Simonin, and Jean P Gibert. Protist predation influences the temperature response of bacterial communities. Frontiers in microbiology, 13, 2022.

31. Yinon M Bar-On, Rob Phillips, and Ron Milo. The biomass distribution on earth. Proceedings of the National Academy of Sciences, 115(25):6506–6511, 2018.

32. Ben Bond-Lamberty and Allison Thomson. Temperature-associated increases in the global soil respiration record. Nature, 464(7288):579–582, 2010.

33. Vincent EJ Jassey, Constant Signarbieux, Stephan Hättenschwiler, Luca Bragazza, Alexandre Buttler, Frédéric Delarue, Bertrand Fournier, Daniel Gilbert, Fatima Laggoun-Défarge, Enrique Lara, et al. An unexpected role for mixotrophs in the response of peatland carbon cycling to climate warming. Scientific reports, 5(1):1–10, 2015.

34. Zhilei Gao, Ida Karlsson, Stefan Geisen, George Kowalchuk, and Alexandre Jousset. Protists: puppet masters of the rhizosphere microbiome. Trends in Plant Science, 24(2): 165–176, 2019.

35. Madhav P Thakur and Stefan Geisen. Trophic regulations of the soil microbiome. Trends in microbiology, 27(9):771–780, 2019.

36. Wu Xiong, Yuqi Song, Keming Yang, Yian Gu, Zhong Wei, George A Kowalchuk, Yangchun Xu, Alexandre Jousset, Qirong Shen, and Stefan Geisen. Rhizosphere protists are key determinants of plant health. Microbiome, 8(1):1–9, 2020.

37. Javier A Ceja-Navarro, Yuan Wang, Daliang Ning, Abelardo Arellano, Leila Ramanculova, Mengting Maggie Yuan, Alyssa Byer, Kelly D Craven, Malay C Saha, Eoin L Brodie, et al. Protist diversity and community complexity in the rhizosphere of switchgrass are dynamic as plants develop. Microbiome, 9(1):1–18, 2021.

38. Daniel Gómez-Pérez, Monja Schmid, Vasvi Chaudhry, Ana Velic, Boris Maček, Ariane Kemen, and Eric Kemen. Proteins released into the plant apoplast by the obligate parasitic protist albugo selectively repress phyllosphere-associated bacteria. bioRxiv, page 11, 2022. doi: 10.1101/2022.05.16.492175.

39. Iqra Bashir, Aadil Farooq War, Iflah Rafiq, Zafar A Reshi, Irfan Rashid, and Yogesh S Shouche. Phyllosphere microbiome: Diversity and functions. Microbiological Research, 254:126888, 2022.

40. Sai Guo, Wu Xiong, Xinnan Hang, Zhilei Gao, Zixuan Jiao, Hongjun Liu, Yani Mo, Nan Zhang, George A Kowalchuk, Rong Li, et al. Protists as main indicators and determinants of plant performance. Microbiome, 9(1):1–11, 2021.

41. James H Brown, James F Gillooly, Andrew P Allen, Van M Savage, and Geoffrey B West. Toward a metabolic theory of ecology. Ecology, 85(7):1771–1789, 2004.

42. D Atkinson. Temperature and organism size: a biological law for ectotherms? Adv Ecol Res, 25:1–58, 1994.

43. Daniel J Wieczynski, Pranav Singla, Adrian Doan, Alexandra Singleton, Ze-Yi Han, Samantha Votzke, Andrea Yammine, and Jean P Gibert. Linking species traits and demography to explain complex temperature responses across levels of organization. Proceedings of the National Academy of Sciences, 118(42), 2021.

44. Jean P Gibert, Rachel L Allen, Ron J Hruska III, and John P DeLong. The ecological consequences of environmentally induced phenotypic changes. Ecology Letters, 20(8): 997–1003, 2017.

45. Douglas S Glazier. A unifying explanation for diverse metabolic scaling in animals and plants. Biological Reviews, 85(1):111–138, 2010.

46. Andrew G Hirst, Douglas S Glazier, and David Atkinson. Body shape shifting during growth permits tests that distinguish between competing geometric theories of metabolic scaling. Ecology Letters, 17(10):1274–1281, 2014.

47. David Atkinson, Benjamin J Ciotti, and David JS Montagnes. Protists decrease in size linearly with temperature: ca. 2.5% c-1. Proceedings of the Royal Society of London. Series B: Biological Sciences, 270(1533):2605–2611, 2003.

48. Jean P Gibert, Ze-Yi Han, Daniel J Wieczynski, Samantha Votzke, and Andrea Yammine. Feedbacks between size and density determine rapid eco-phenotypic dynamics. Functional Ecology, 36(7):1668–1680, 2022.

49. Y Nakano, K Miyatake, H Okuno, K Hamazaki, S Takenaka, N Honami, M Kiyota, I Aiga, and J Kondo. Growth of photosynthetic algae euglena in high co2 conditions and its photosynthetic characteristics. In International Symposium on Plant Production in Closed Ecosystems 440, pages 49–54, 1996.

50. Monika K Reczuga, Christophe Victor William Seppey, Matthieu Mulot, Vincent EJ Jassey, Alexandre Buttler, Sandra Słowińska, Michał Słowiński, Enrique Lara, Mariusz Lamentowicz, and Edward AD Mitchell. Assessing the responses of sphagnum micro-eukaryotes to climate changes using high throughput sequencing. PeerJ, 8:e9821, 2020.

51. Andrey N Tsyganov, Rien Aerts, Ivan Nijs, Johannes HC Cornelissen, and Louis Beyens. Sphagnum-dwelling testate amoebae in subarctic bogs are more sensitive to soil warming in the growing season than in winter: the results of eight-year field climate manipulations. Protist, 163(3):400–414, 2012.

52. Vincent EJ Jassey, Mariusz Lamentowicz, Luca Bragazza, Maaike L Hofsommer, Robert TE Mills, Alexandre Buttler, Constant Signarbieux, and Bjorn JM Robroek. Loss of testate amoeba functional diversity with increasing frost intensity across a continental gradient reduces microbial activity in peatlands. European journal of protistology, 55: 190–202, 2016.

53. Anna M Basińska, Monika K Reczuga, Maciej Gąbka, Marcin Stróżecki, Dominika Łuców, Mateusz Samson, Marek Urbaniak, Jacek Lesńy, Bogdan H Chojnicki, Daniel Gilbert, et al. Experimental warming and preci ? pitation reduction affect the biomass of microbial communities in a sphagnum peatland. Ecological Indicators, 112:106059, 2020.

54. Cyrille Violle, Marie-Laure Navas, Denis Vile, Elena Kazakou, Claire Fortunel, Irène Hummel, and Eric Garnier. Let the concept of trait be functional! Oikos, 116(5):882–892, 2007.

55. David JS Montagnes, Susan A Kimmance, and David Wilson. Effects of global and local temperature changes on free living, aquatic protists. In Conference Proceedings of the 14th International Conference on Comparative Physiology, Climate Changes: Effects on Plants, Animals, and Humans (eds Bolis CL, Keines R, Elia M et al.). Available on CD ROM (from the senior author), pages 1–13, 2002.

56. A Clarke and KPP Fraser. Why does metabolism scale with temperature? Functional ecology, 18(2):243–251, 2004.

57. K. E. Fussmann, B. Rosenbaum, U. Brose, and B.C. Rall. Interactive effects of shifting body size and feeding adaptation drive interaction strengths of protist predators under warming. bioRxiv, pages 1–34, 2017. doi: 10.1101/101675.

58. Susanne Menden-Deuer, Caitlyn Lawrence, and Gayantonia Franzè. Herbivorous protist growth and grazing rates at in situ and artificially elevated temperatures during an arctic phytoplankton spring bloom. PeerJ, 6:e5264, 2018.

59. Gabriel Yvon-Durocher, J Iwan Jones, Mark Trimmer, Guy Woodward, and Jose M Montoya. Warming alters the metabolic balance of ecosystems. Philosophical Transactions of the Royal Society B: Biological Sciences, 365(1549):2117–2126, 2010.

60. Susanne Wilken, Jef Huisman, Suzanne Naus-Wiezer, and Ellen Van Donk. Mixotrophic organisms become more heterotrophic with rising temperature. Ecology letters, 16(2): 225–233, 2013.

61. M Matsubayashi, H Ando, I Kimata, H Nakagawa, M Furuya, H Tani, and K Sasai. Morphological changes and viability of cryptosporidium parvum sporozoites after excystation in cell-free culture media. Parasitology, 137(13):1861–1866, 2010.

62. AS Sediq, R Klem, MR Nejadnik, P Meij, and Wim Jiskoot. Label-free, flow-imaging methods for determination of cell concentration and viability. Pharmaceutical research, 35(8): 1–10, 2018.

63. Ulrich Brose, Jennifer A Dunne, Jose M Montoya, Owen L Petchey, Florian D Schneider, and Ute Jacob. Climate change in size-structured ecosystems, 2012.

64. Jean P Gibert. Temperature directly and indirectly influences food web structure. Scientific reports, 9(1):5312, 2019.

65. Mary I O’Connor, Michael F Piehler, Dina M Leech, Andrea Anton, and John F Bruno. Warming and resource availability shift food web structure and metabolism. PLoS biology, 7(8):e1000178, 2009.

66. Daniel J Wieczynski, Kristin M Yoshimura, Elizabeth R Denison, Stefan Geisen, Jennifer M DeBruyn, A Jonathan Shaw, David J Weston, Dale A Pelletier, Steven W Wilhelm, and Jean P Gibert. Viral infections likely mediate microbial controls on ecosystem responses to global warming. FEMS Microbiology Ecology, page fiad016, 2023.

67. Paul J Hanson, Jeffery S Riggs, W Robert Nettles, Jana R Phillips, Misha B Krassovski, Leslie A Hook, Lianhong Gu, Andrew D Richardson, Donald M Aubrecht, Daniel M Ricciuto, et al. Attaining whole-ecosystem warming using air and deep-soil heating methods with an elevated co 2 atmosphere. Biogeosciences, 14(4):861–883, 2017.

68. David Atkinson, Simon A Morley, and Roger N Hughes. From cells to colonies: at what levels of body organization does the ‘temperature-size rule’apply? Evolution & development, 8(2):202–214, 2006.

69. HyeonSeok Shin, Seong-Joo Hong, Chan Yoo, Mi Han, Hookeun Lee, Hyung-Kyoon Choi, Suhyung Cho, Choul-Gyun Lee, and Byung-Kwan Cho. Genome-wide transcriptome analysis revealed organelle specific responses to temperature variations in algae. Scientific reports, 6(1):1–11, 2016.

70. Ulrich Sommer, Carolin Paul, and Maria Moustaka-Gouni. Warming and ocean acidification effects on phytoplankton—from species shifts to size shifts within species in a mesocosm experiment. PloS one, 10(5):e0125239, 2015.

71. Ulrich Sommer, Kalista H Peter, Savvas Genitsaris, and Maria Moustaka-Gouni. Do marine phytoplankton follow b ergmann’s rule sensu lato? Biological Reviews, 92(2):1011–1026, 2017.

72. PJ Hanson, AL Gill, X Xu, JR Phillips, DJ Weston, RK Kolka, JS Riggs, and LA Hook. Intermediate-scale community-level flux of co2 and ch4 in a minnesota peatland: putting the spruce project in a global context. Biogeochemistry, 129(3):255–272, 2016.

73. Malak M Tfaily, William T Cooper, Joel E Kostka, Patrick R Chanton, Christopher W Schadt, Paul J Hanson, Colleen M Iversen, and Jeffrey P Chanton. Organic matter transformation in the peat column at marcell experimental forest: humification and vertical stratification. Journal of Geophysical Research: Biogeosciences, 119(4):661–675, 2014.

74. Diane K Stoecker, Dian J Gifford, and Mary Putt. Preservation of marine planktonic ciliates: losses and cell shrinkage during fixation. Marine Ecology Progress Series, pages 293–299, 1994.

75. Monica Modigh and Sara Castaldo. Effects of fixatives on ciliates as related to cell size. Journal of Plankton Research, 27(8):845–849, 2005.

76. Chris Fraley and Adrian E Raftery. Model-based clustering, discriminant analysis, and density estimation. Journal of the American statistical Association, 97(458):611–631, 2002.

77. Alboukadel Kassambara and Maintainer Alboukadel Kassambara. Package ‘ggpubr’. R package version 0.1, 6, 2020.

78. Simon Wood and Maintainer Simon Wood. Package ‘mgcv’. R package version, 1(29): 729, 2015.

79. Yves Rosseel. lavaan: An r package for structural equation modeling. Journal of statistical software, 48:1–36, 2012.

80. Etienne Laliberté, Pierre Legendre, Bill Shipley, and Maintainer Etienne Laliberté. Package ‘fd’. Measuring functional diversity from multiple traits, and other tools for functional ecology, 1:0–12, 2014.

81. Matthias Grenié and Hugo Gruson. fundiversity: a modular r package to compute functional diversity indices. Ecography, 2023(3):e06585, 2023.

82. Luisa W Hugerth, Emilie EL Muller, Yue OO Hu, Laura AM Lebrun, Hugo Roume, Daniel Lundin, Paul Wilmes, and Anders F Andersson. Systematic design of 18s rrna gene primers for determining eukaryotic diversity in microbial consortia. PloS one, 9(4):e95567, 2014.

83. Benjamin J Callahan, Paul J McMurdie, Michael J Rosen, Andrew W Han, Amy Jo A Johnson, and Susan P Holmes. Dada2: High-resolution sample inference from illumina amplicon data. Nature methods, 13(7):581–583, 2016.

84. Laure Guillou, Dipankar Bachar, Stéphane Audic, David Bass, Cédric Berney, Lucie Bittner, Christophe Boutte, Gaétan Burgaud, Colomban de Vargas, Johan Decelle, et al. The protist ribosomal reference database (pr2): a catalog of unicellular eukaryote small subunit rrna sequences with curated taxonomy. Nucleic acids research, 41(D1):D597–D604, 2012.

85. Paul J McMurdie and Susan Holmes. phyloseq: an r package for reproducible interactive analysis and graphics of microbiome census data. PloS one, 8(4):e61217, 2013.

86. Roberto Danovaro, Antonio Dell’Anno, and Antonio Pusceddu. Biodiversity response to climate change in a warm deep sea. Ecology Letters, 7(9):821–828, 2004.

87. Luke O Frishkoff, Daniel S Karp, Leithen K M’Gonigle, Chase D Mendenhall, Jim Zook, Claire Kremen, Elizabeth A Hadly, and Gretchen C Daily. Loss of avian phylogenetic diversity in neotropical agricultural systems. science, 345(6202):1343–1346, 2014.

88. Zhenghu Zhou, Chuankuan Wang, and Yiqi Luo. Meta-analysis of the impacts of global change factors on soil microbial diversity and functionality. Nature communications, 11(1): 1–10, 2020.

89. Håkan Rydin, Urban Gunnarsson, and Sebastian Sundberg. The role of sphagnum in peatland development and persistence. In Boreal peatland ecosystems, pages 47–65. Springer, 2006.

90. JM Waddington, M Strack, and MJ Greenwood. Toward restoring the net carbon sink function of degraded peatlands: Short-term response in co2 exchange to ecosystem-scale restoration. Journal of Geophysical Research: Biogeosciences, 115(G1), 2010.

91. Richard D Bardgett, Chris Freeman, and Nicholas J Ostle. Microbial contributions to climate change through carbon cycle feedbacks. The ISME journal, 2(8):805–814, 2008.

92. Jizhong Zhou, Kai Xue, Jianping Xie, YE Deng, Liyou Wu, Xiaoli Cheng, Shenfeng Fei, Shiping Deng, Zhili He, Joy D Van Nostrand, et al. Microbial mediation of carbon-cycle feedbacks to climate warming. Nature Climate Change, 2(2):106–110, 2012.

93. Thomas P Smith, Thomas JH Thomas, Bernardo García-Carreras, Sofía Sal, Gabriel Yvon-Durocher, Thomas Bell, and Samrāt Pawar. Community-level respiration of prokaryotic microbes may rise with global warming. Nature communications, 10(1):1–11, 2019.

94. James F Gillooly, James H Brown, Geoffrey B West, Van M Savage, and Eric L Charnov. Effects of size and temperature on metabolic rate. science, 293(5538):2248–2251, 2001.

95. Janet L Gardner, Anne Peters, Michael R Kearney, Leo Joseph, and Robert Heinsohn. Declining body size: a third universal response to warming? Trends in ecology & evolution, 26(6):285–291, 2011.

96. Jean P Gibert and John P DeLong. Temperature alters food web body-size structure. Biology letters, 10(8):20140473, 2014.

97. Wilco CEP Verberk, David Atkinson, K Natan Hoefnagel, Andrew G Hirst, Curtis R Horne, and Henk Siepel. Shrinking body sizes in response to warming: explanations for the temperature–size rule with special emphasis on the role of oxygen. Biological Reviews, 96 (1):247–268, 2021.

98. John P DeLong, Chad E Brassil, Emma K Erickson, Valery E Forbes, Etsuko N Moriyama, and Wayne R Riekhof. Dynamic thermal reaction norms and body size oscillations challenge explanations of the temperature–size rule. Evolutionary Ecology Research, 18(3): 293–303, 2017.

99. John P DeLong, Jordan G Okie, Melanie E Moses, Richard M Sibly, and James H Brown. Shifts in metabolic scaling, production, and efficiency across major evolutionary transitions of life. Proceedings of the National Academy of Sciences, 107(29):12941–12945, 2010.

100. Jean P Gibert, Daniel J Wieczynski, Ze-Yi Han, and Andrea Yammine. Rapid ecophenotypic feedback and the temperature response of biomass dynamics. Ecology and Evolution, 13(1):e9685, 2023.

101. Coral J Fung Shek. Dynamic Imaging Particle Technology for Quantitative Morphological Analysis and Cell Counting. PhD thesis, Johns Hopkins University, 2015.

102. Samuel Hamard, Regis Céréghino, Maialen Barret, Anna Sytiuk, Enrique Lara, Ellen Dorrepaal, Paul Kardol, Martin Küttim, Mariusz Lamentowicz, Joséphine Leflaive, et al. Contribution of microbial photosynthesis to peatland carbon uptake along a latitudinal gradient. Journal of Ecology, 109(9):3424–3441, 2021.

103. Vincent EJ Jassey, Samuel Hamard, Cécile Lepère, Régis Céréghino, Bruno Corbara, Martin Küttim, Joséphine Leflaive, Céline Leroy, and Jean-François Carrias. Photosynthetic microorganisms effectively contribute to bryophyte co2 fixation in boreal and tropical regions. ISME Communications, 2(1):64, 2022.

104. Vincent EJ Jassey, Romain Walcker, Paul Kardol, Stefan Geisen, Thierry Heger, Mariusz Lamentowicz, Samuel Hamard, and Enrique Lara. Contribution of soil algae to the global carbon cycle. New Phytologist, 234(1):64–76, 2022.

105. Lennart T Bach, Santiago Alvarez-Fernandez, Thomas Hornick, Annegret Stuhr, and Ulf Riebesell. Simulated ocean acidification reveals winners and losers in coastal phytoplankton. PloS one, 12(11):e0188198, 2017.

106. Alexandra Coello-Camba, Susana Agustí, Johnna Holding, Jesús M Arrieta, and Carlos M Duarte. Interactive effect of temperature and co2 increase in arctic phytoplankton. Frontiers in Marine Science, 1:49, 2014.

107. Andrew Clarke. Principles of thermal ecology: Temperature, energy and life. Oxford University Press, 2017.

108. Allison R Rober, Kevin S McCann, Merritt R Turetsky, and Kevin H Wyatt. Cascading effects of predators on algal size structure. Journal of Phycology, 58(2):308–317, 2022.

109. Edward B Rastetter, Göran I Ågren, and Gaius R Shaver. Responses of n-limited ecosystems to increased co2: A balanced-nutrition, coupled-element-cycles model. Ecological Applications, 7(2):444–460, 1997.

110. Petra MA Fransson and Emma M Johansson. Elevated co2 and nitrogen influence exudation of soluble organic compounds by ectomycorrhizal root systems. FEMS Microbiology Ecology, 71(2):186–196, 2010.

111. Edward AD Mitchell, Daniel Gilbert, Alexandre Buttler, Christian Amblard, Philippe Grosvernier, and J-M Gobat. Structure of microbial communities in sphagnum peatlands and effect of atmospheric carbon dioxide enrichment. Microbial ecology, 46(2):187–199, 2003.

112. Caitlin Petro, Alyssa A Carrell, Rachel M Wilson, Katherine Duchesneau, Sekou Noble-Kuchera, Tianze Song, Colleen M Iversen, Joanne Childs, Geoff Schwaner, Jeffrey P Chanton, et al. Climate drivers alter nitrogen availability in surface peat and decouple n2 fixation from ch4 oxidation in the sphagnum moss microbiome. Global Change Biology, 2023.

113. Luca Bragazza, Julien Parisod, Alexandre Buttler, and Richard D Bardgett. Biogeochemical plant–soil microbe feedback in response to climate warming in peatlands. Nature Climate Change, 3(3):273–277, 2013.

114. Kathrin Rousk. Biotic and abiotic controls of nitrogen fixation in cyanobacteria–moss associations. New Phytologist, 235(4):1330–1335, 2022.

115. Danillo O Alvarenga and Kathrin Rousk. Indirect effects of climate change inhibit n2 fixation associated with the feathermoss hylocomium splendens in subarctic tundra. Science of the Total Environment, 795:148676, 2021.

116. Ina C Meier, Seth G Pritchard, Edward R Brzostek, M Luke McCormack, and Richard P Phillips. The rhizosphere and hyphosphere differ in their impacts on carbon and nitrogen cycling in forests exposed to elevated co 2. New Phytologist, 205(3):1164–1174, 2015.

117. Shauna M Uselman, Robert G Qualls, and Richard B Thomas. A test of a potential short cut in the nitrogen cycle: The role of exudation of symbiotically fixed nitrogen from the roots of a n-fixing tree and the effects of increased atmospheric co2 and temperature. Plant and Soil, 210(1):21–32, 1999.

118. Takehito Yoshida, Laura E Jones, Stephen P Ellner, Gregor F Fussmann, and Nelson G Hairston. Rapid evolution drives ecological dynamics in a predator–prey system. Nature, 424(6946):303–306, 2003.

119. Ze-Yi Han, Daniel J Wieczynski, Andrea Yammine, and Jean P Gibert. Temperature and nutrients drive eco-phenotypic dynamics in a microbial food web. Proceedings of the Royal Society B, 290(1992):20222263, 2023.

120. John P DeLong and Jean P Gibert. Gillespie eco-evolutionary models (gem s) reveal the role of heritable trait variation in eco-evolutionary dynamics. Ecology and evolution, 6(4): 935–945, 2016.

121. Thomas M Luhring and John P DeLong. Trophic cascades alter eco-evolutionary dynamics and body size evolution. Proceedings of the Royal Society B, 287(1938):20200526, 2020.

122. Matthew J Amesbury, Angela Gallego-Sala, and Julie Loisel. Peatlands as prolific carbon sinks. Nature Geoscience, 12(11):880–881, 2019.

123. Angela M Oliverio, Stefan Geisen, Manuel Delgado-Baquerizo, Fernando T Maestre, Benjamin L Turner, and Noah Fierer. The global-scale distributions of soil protists and their contributions to belowground systems. Science advances, 6(4):eaax8787, 2020.

124. Mara Y McPartland, Evan S Kane, Michael J Falkowski, Randy Kolka, Merritt R Turetsky, Brian Palik, and Rebecca A Montgomery. The response of boreal peatland community composition and ndvi to hydrologic change, warming, and elevated carbon dioxide. Global change biology, 25(1):93–107, 2019.

125. Cara A Faillace, Arnaud Sentis, and José M Montoya. Eco-evolutionary consequences of habitat warming and fragmentation in communities. Biological Reviews, 96(5):1933–1950, 2021.

126. Marianne Simon, Purificación López-García, Philippe Deschamps, David Moreira, Gwendal Restoux, Paola Bertolino, and Ludwig Jardillier. Marked seasonality and high spatial variability of protist communities in shallow freshwater systems. The ISME journal, 9(9): 1941–1953, 2015.

127. Johan De Gruyter, James T Weedon, Stéphane Bazot, Steven Dauwe, Pere-Roc Fernandez-Garberí, Stefan Geisen, Louis Gourlez De La Motte, Bernard Heinesch, Ivan A Janssens, Niki Leblans, et al. Patterns of local, intercontinental and interseasonal variation of soil bacterial and eukaryotic microbial communities. FEMS microbiology ecology, 96(3): fiaa018, 2020.

128. Julia Schroeder, Lisa Kammann, Mirjam Helfrich, Christoph C Tebbe, and Christopher Poeplau. Impact of common sample pre-treatments on key soil microbial properties. Soil Biology and Biochemistry, 160:108321, 2021.

129. Vincent EJ Jassey, Geneviève Chiapusio, Philippe Binet, Alexandre Buttler, Fatima Laggoun-Défarge, Frédéric Delarue, Nadine Bernard, Edward AD Mitchell, Marie-Laure Toussaint, André-Jean Francez, et al. Above-and belowground linkages in sphagnum peatland: Climate warming affects plant-microbial interactions. Global Change Biology, 19(3):811–823, 2013.

130. Xiao-Ying Ma, Hao Xu, Zi-Yin Cao, Lei Shu, and Rui-Liang Zhu. Will climate change cause the global peatland to expand or contract? evidence from the habitat shift pattern of sphagnum mosses. Global Change Biology, 28(21):6419–6432, 2022.

131. Aditee Mitra, Kevin J Flynn, Joann M Burkholder, Terrje Berge, Albert Calbet, John A Raven, Edna Granéli, Pat M Glibert, Per Juel Hansen, Diane K Stoecker, et al. The role of mixotrophic protists in the biological carbon pump. Biogeosciences, 11(4):995–1005, 2014.

132. Zhenfeng Liu, Victoria Campbell, Karla B Heidelberg, and David A Caron. Gene expression characterizes different nutritional strategies among three mixotrophic protists. FEMS Microbiology Ecology, 92(7):fiw106, 2016.

133. Marc-André Selosse, Marie Charpin, and Fabrice Not. Mixotrophy everywhere on land and in water: the grand écart hypothesis. Ecology Letters, 20(2):246–263, 2017.

134. Daniel Wieczynski, Holly Moeller, and Jean-Philippe Gibert. Mixotrophs generate carbon tipping points under warming, 2022.

135. J Limpens, F Berendse, Christian Blodau, JG Canadell, C Freeman, J Holden, N Roulet, Håkan Rydin, and Gabriela Schaepman-Strub. Peatlands and the carbon cycle: from local processes to global implications–a synthesis. Biogeosciences, 5(5):1475–1491, 2008.

136. Steve Frolking, Julie Talbot, Miriam C Jones, Claire C Treat, J Boone Kauffman, Eeva-Stiina Tuittila, and Nigel Roulet. Peatlands in the earth’s 21st century climate system. Environmental Reviews, 19(NA):371–396, 2011.

137. Eville Gorham, Clarence Lehman, Arthur Dyke, Dicky Clymo, and Joannes Janssens. Long-term carbon sequestration in north american peatlands. Quaternary Science Reviews, 58:77–82, 2012.

138. Qinglong You, Ziyi Cai, Nick Pepin, Deliang Chen, Bodo Ahrens, Zhihong Jiang, Fangying Wu, Shichang Kang, Ruonan Zhang, Tonghua Wu, et al. Warming amplification over the arctic pole and third pole: Trends, mechanisms and consequences. Earth-Science Reviews, 217:103625, 2021.

