## Supplemental Materials for "Climate Change Factors Interactively Shift Peatland Functional Microbial Composition in a Whole-Ecosystem Warming Experiment"

### 1: Supplemental Materials

**A. Materials and Methods.** We obtained 37 physical and optical functional composition traits from the image libraries (Table S1, Fig. S1A). Initial exploratory data analysis indicated that many of these traits were skewed and we subsequently applied transformations to normalize the data (Table S1). Following normalization, we selected a subset of traits to model across-temperature and carbon dioxide treatments by eliminating highly-correlated traits (Fig. S1) within a subset of our functional composition analyses (i.e. size, shape, contents, and resource acquisition). We chose these functional trait subsets from first principles of metabolic theory and ecology (1). By choosing one trait within each subset, we reduced co-linearity (Fig. S1B), which also increased modeling power. The final measurements included volume (organismal size), aspect ratio (cell shape), sigma intensity (cellular contents), and red/green ratio (resource acquisition mode).

We calculated Geodesic Length and Thickness trait values assuming the cellular particle is modeled as a rectangle, with perimeter ($P$) defined as the length of a particle edge excluding the observed lengths of the edges of holes in said particle (Cite FlowCam literature manual). To calculate physical traits, we assumed the protist cell to be a prolate spheroid, with volume (V) and aspect ratio dependent upon the geodesic width (W, shortest axis) and geodesic length (L, longest axis) of each cell, such that V = 4$\pi$W23L. We then define aspect ratio (AR) as the ratio of geodesic width to geodesic length AR = WL. Due to shadows and other optical artifacts, the FlowCam software occasionally assigns protist cells an erroneous aspect ratio value of 1 (e.g. perfect spheroid); we removed these observations from our analysis, leading to a final data set size of n = 157,177 observations. Optical traits were derived from the FlowCam systems proprietary software, with sigma intensity defined as the standard deviation of the grayscale values of a given cell. And, the red/green ratio obtained by comparing the pixel intensity of the red channel and green channel with a background of the images' total reflective intensity. From this assumption, the physical and optical traits for individual cells were calculated:

**Geodesic Length (*L*)**

$$L = \frac{P + \sqrt{P^2 - 16 \times (\frac{\pi \times diameter^2}{4})}}{4}$$

**Geodesic Thickness (*W*)**

$$W = \frac{P - \sqrt{P^2 - 16 \times (\frac{\pi \times diameter^2}{4})}}{4}$$

**Geodesic Aspect Ratio (*AR*)**

$$AR = \frac{W}{L}$$

**Volume (*V*)**

$$V = \frac{4\pi}{3} \times \frac{(\frac{W}{2})^2}{(\frac{L}{2})}$$

**Sigma Intensity (*S*)**

$$S = \sqrt{\frac{(\sum_{i=1}^{n}\sum_{i=1}^{m} x_{i,j}^2 \times [y_{i,j} = 1]) - \frac{SumIntensity^2}{pixelcount}}{pixelcount}}$$

where,

$$pixelcount = (\sum_{i=1}^{n}\sum_{i=1}^{m}[y_{i,j} = 1])$$

and,

$$sumintensity = (\sum_{i=1}^{n}\sum_{i=1}^{m} x_{i,j} \times [y_{i,j} = 1])$$

when *n,m* are the height and width of the image; *i,j* are row and column indices; $x_{i,j}$ is a pixel intensity in the grayscale image; and $y_{i,j}$ is a pixel intensity in the binary image.

**Average Red to Average Green Ratio (*R/G*)**

$$R/G = \frac{AverageRed}{AverageGreen}$$

where,

$$AverageRed = \frac{(\sum_{i=1}^{n}\sum_{i=1}^{m} x_{i,j} \times [y_{i,j} = 1])}{(\sum_{i=1}^{n}\sum_{i=1}^{m}[y_{i,j} = 1])}$$

with *n,m* as the height and width of the image; *i,j* as row and column indices; $x_{i,j}$ as pixel intensity in the red wavelength of the image; and $y_{i,j}$ as the pixel intensity in the binary image. And,

$$AverageGreen = \frac{(\sum_{i=1}^{n}\sum_{i=1}^{m} x_{i,j} \times [y_{i,j} = 1])}{(\sum_{i=1}^{n}\sum_{i=1}^{m}[y_{i,j} = 1])}$$

with *n,m* as the height and width of the image; *i,j* as row and column indices; $x_{i,j}$ as pixel intensity in the green wavelength of the image; and $y_{i,j}$ as the pixel intensity in the binary image.

Due to inefficiencies in capturing all individual cells through fluid imaging as the density of particles increases, we sought to properly estimate the abundance of protists within each treatment, such that the estimated count regressed with the particle density (D) estimation provided by the FlowCam proprietary software creates a 1-to-1 line (Fig. S2); accordingly, $\hat{E}$ = Em, where E is the observed count and m is the regression slope forced through the origin: f (E) = m*D + 0.

**B. Citations.**


1. Daniel J Wieczynski, Pranav Singla, Adrian Doan, Alexandra Singleton, Ze-Yi Han, Samantha Votzke, Andrea Yammine, and Jean P Gibert. Linking species traits and demography to explain complex temperature responses across levels of organization. *Proceedings of the National Academy of Sciences*, 118(42), 2021.
2. Luca Scrucca, Michael Fop, T Brendan Murphy, and Adrian E Raftery. mclust 5: clustering, classification and density estimation using gaussian finite mixture models. *The R journal*, 8 (1):289, 2016.


DRAFT

#### C. Figures.

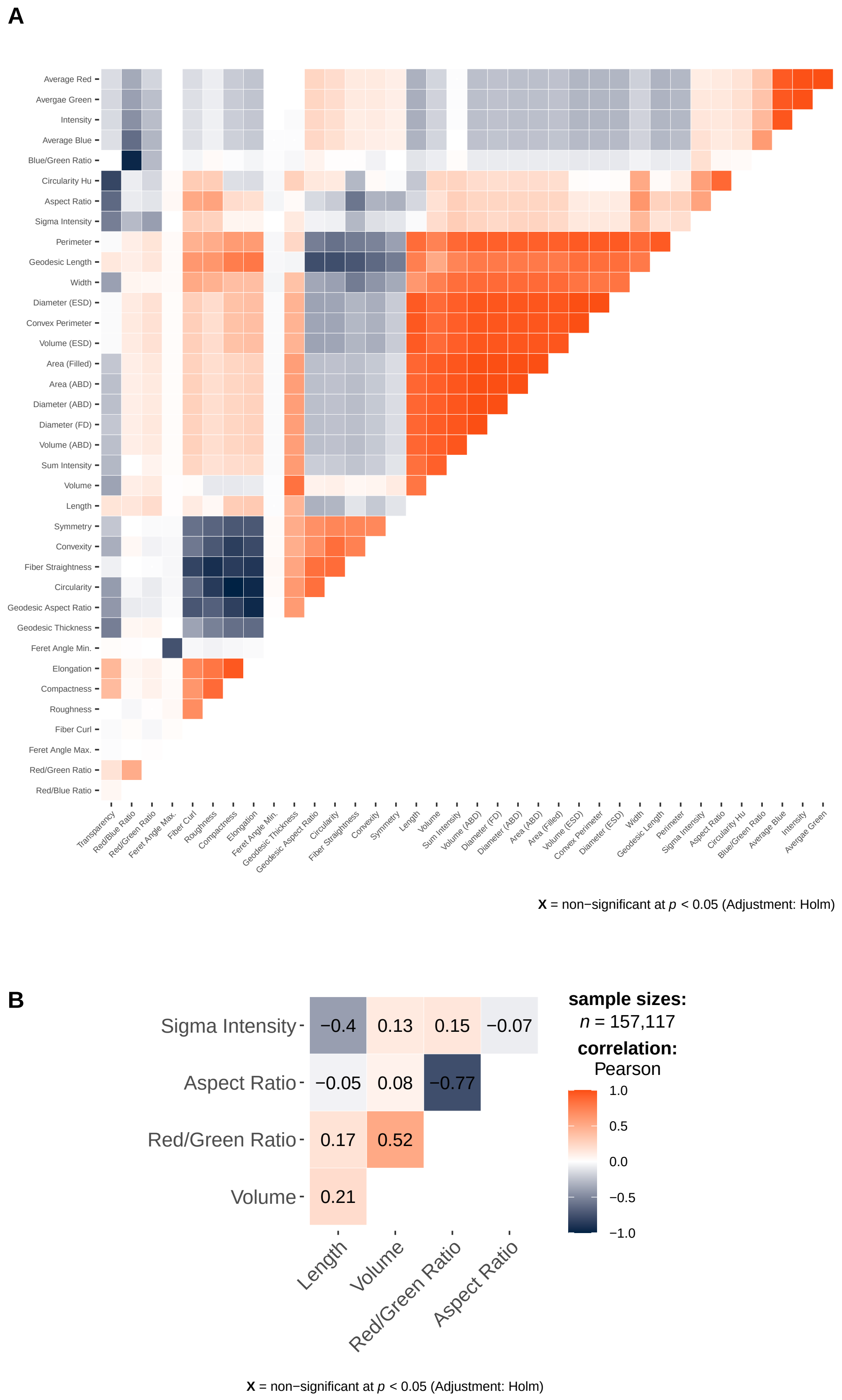


**Fig. S1.** Correlations of functional composition (traits) for (A) all traits measured via fluid imaging and those selected from (B) subsets of trait types (size, shape, cellular contents, and resource acquisition). Pearson correlation value labeled in (B), with color ranges in (A) and (B). All correlations significant in B.

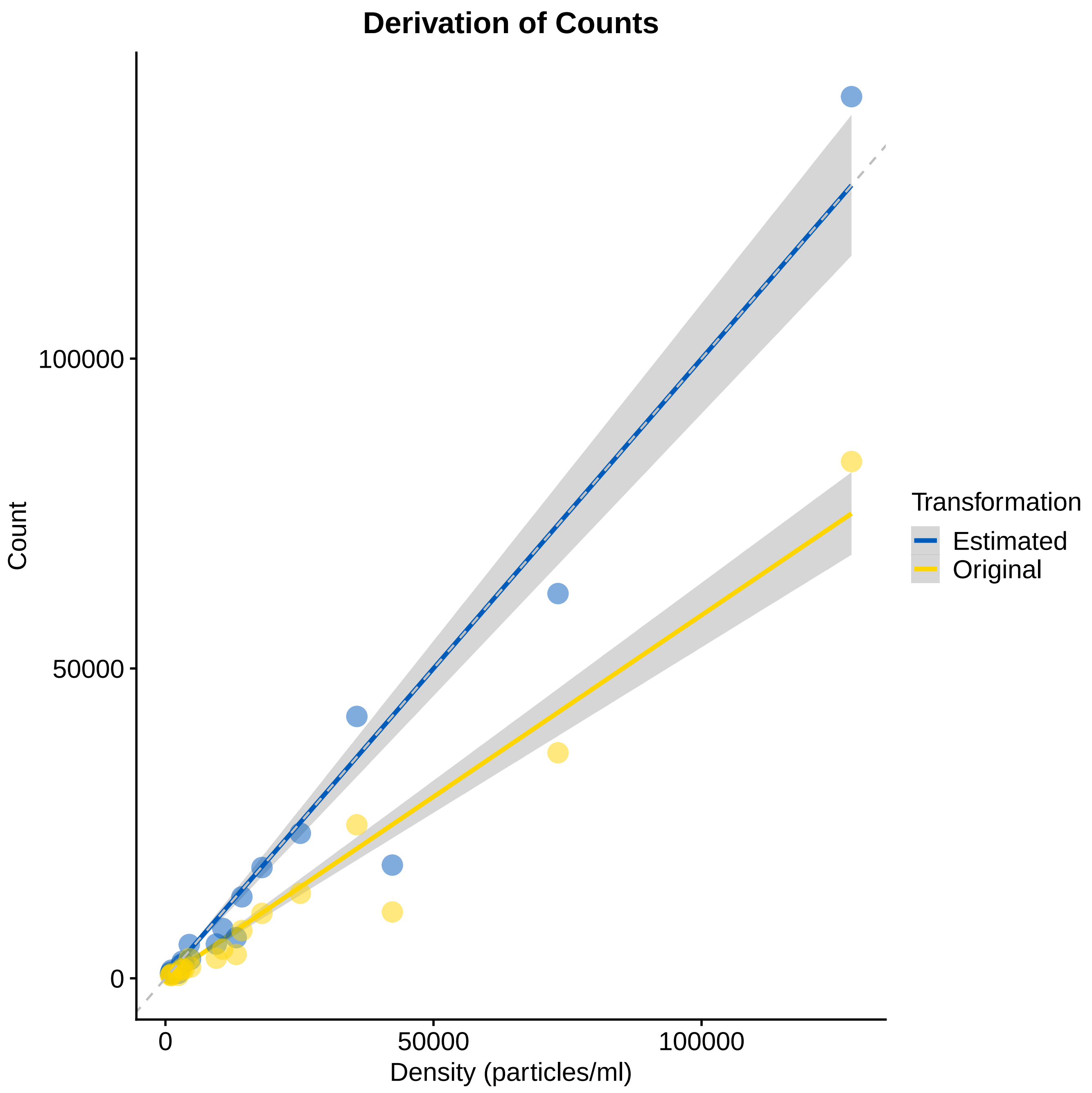


**Fig. S2.** Transformation of raw abundance counts (yellow) to estimated true abundances (blue) based on density counts of the FlowCam. The efficiency of fluid imaging decreases in accurate counting of particles as density of particles increases (yellow slope $\leq 1$). By dividing the raw counts (yellow) by the mean slope, we can adjust the count estimates to obtain a slope = 1 (blue, i.e. observed counts = density of counts).

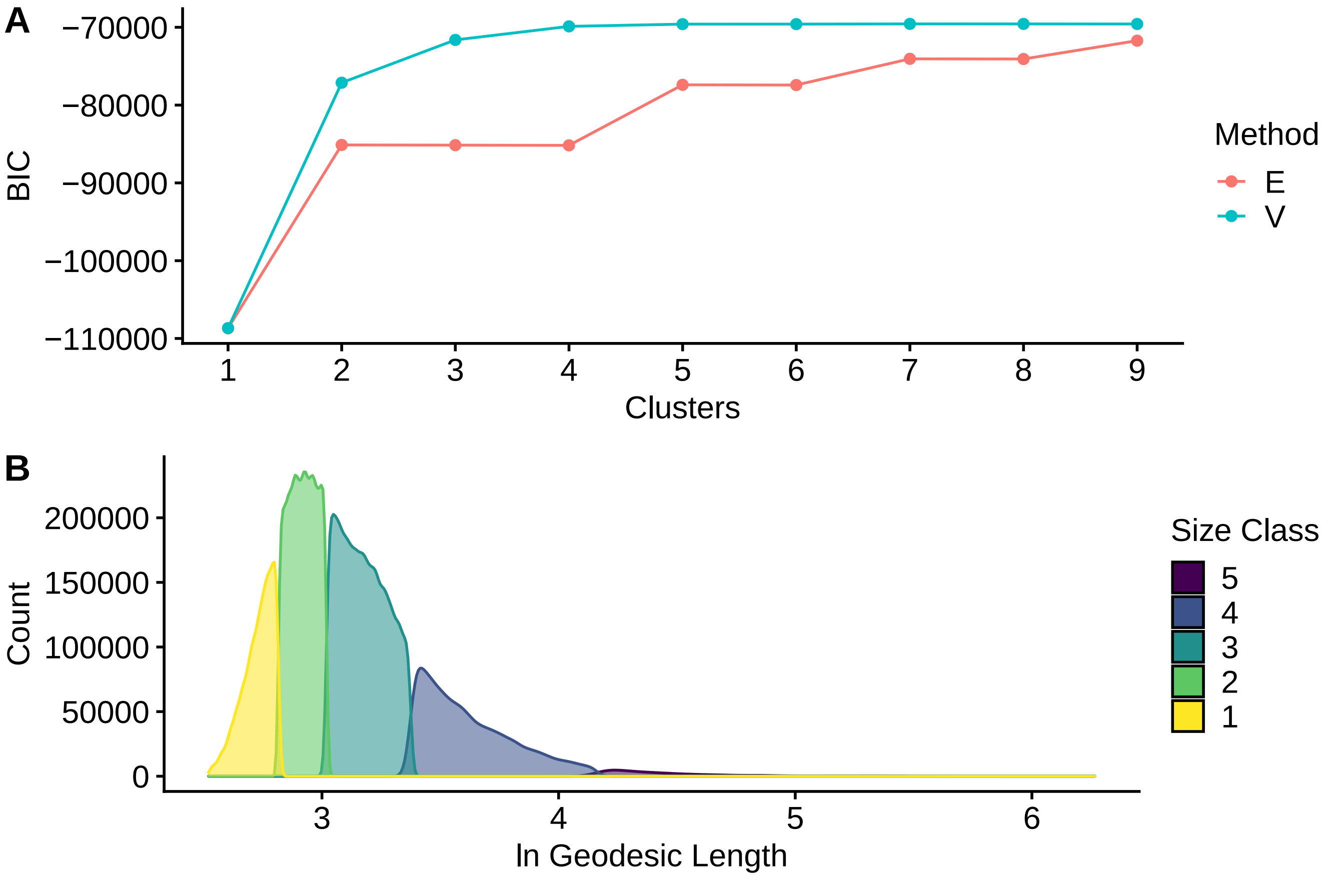


**Fig. S3.** Clustering selection. (A) Bayesian Information Criterion (BIC) by model type (E: Equal Variance, V: Varying Variance) and number of clusters (2). Less negative score indicates higher parsimony—optimal at 5 clusters with V algorithm. (B) Density of clustered size classes generated from a volume-based Bayesian clustering algorithm performed on the geodesic length of individual protists. Geodesic length ranges of exponentiated size class delineations as follows. Size Class 1: 12.43 $\mu$m - 17.07 $\mu$m; Size Class 2: 17.07 $\mu$m - 20.68 $\mu$m; Size Class 3: 20.68 $\mu$m - 28.24 $\mu$m; Size Class 4: 28.24 $\mu$m - 59.74$\mu$m; Size Class 5: 59.74 $\mu$m - 526.43 $\mu$m

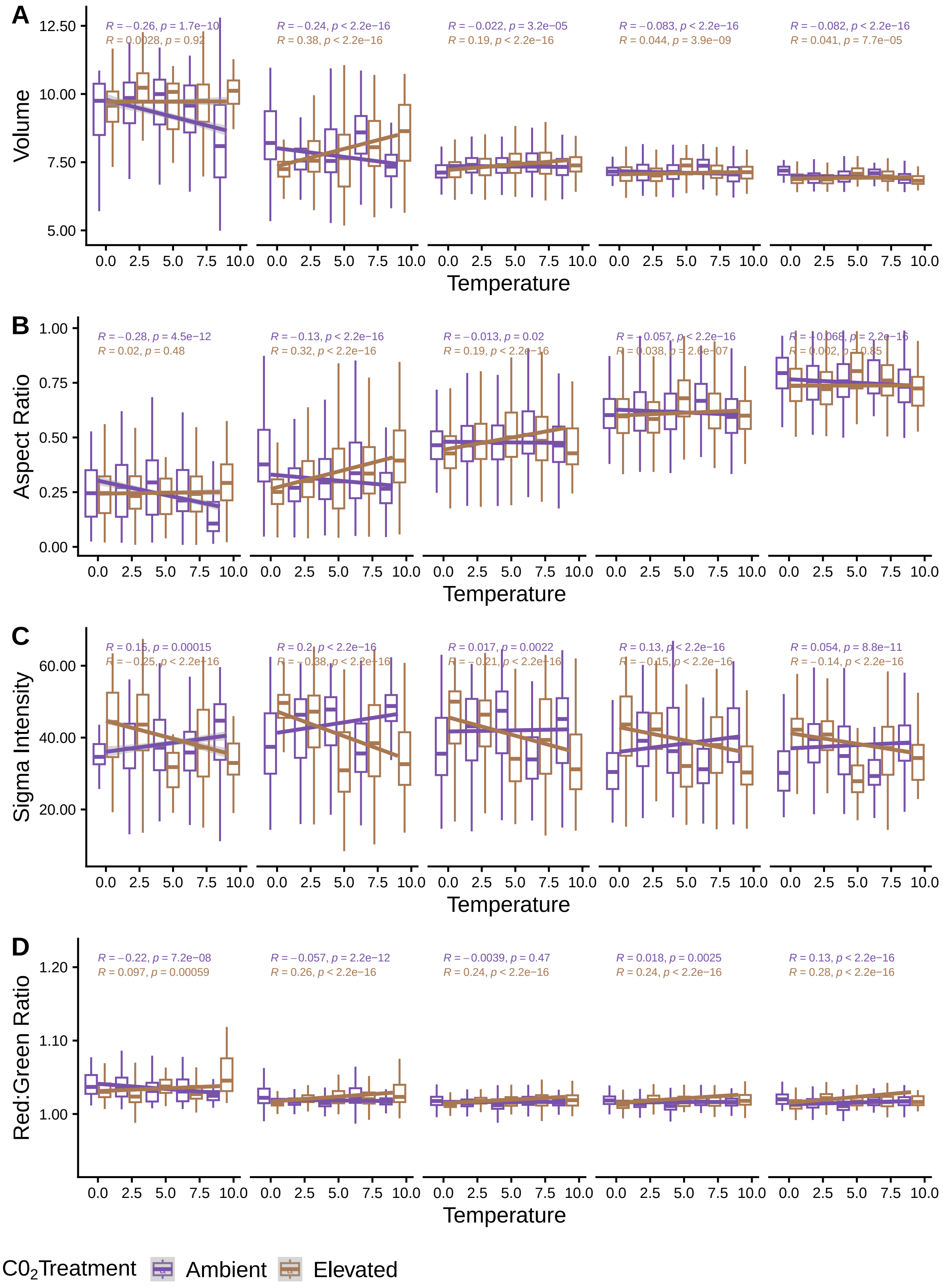


**Fig. S4.** Three-way interaction of functional traits. Significant empirical linear regressions for ambient $CO_2$ (purple lines) and elevated $CO_2$ (gold lines) across temperatures. Generlized linear model (GLM) regression bound by 95% confidence intervals (light gray). All relationships were highly robust and significant $(2.2x10^{-16}) \leq$ p $\leq$ 0.019) across all factorial treatments. Volume (A) shows an increase across temperatures under elevated $CO_2$ amplified by size class and a decrease across temperatures under ambient $CO_2$ also accelerated by size class. Aspect ratio (B) shows the same pattern as volume (A), with a three-way interaction. Sigma intensity (C) displays an inverse pattern to volume (A) and aspect ratio (B), with larger size classes amplifying an increase in cellular contents under ambient $CO_2$ and a decrease under elevated $CO_2$. Finally, the reg/green ratio (D) regression indicates a three-way interaction, with less clear of a pattern. While elevated $CO_2$ consistently leads to redder cells augmented by size class, ambient $CO_2$ induces a reversal in trends for the three largest size classes, which get increasingly greener.

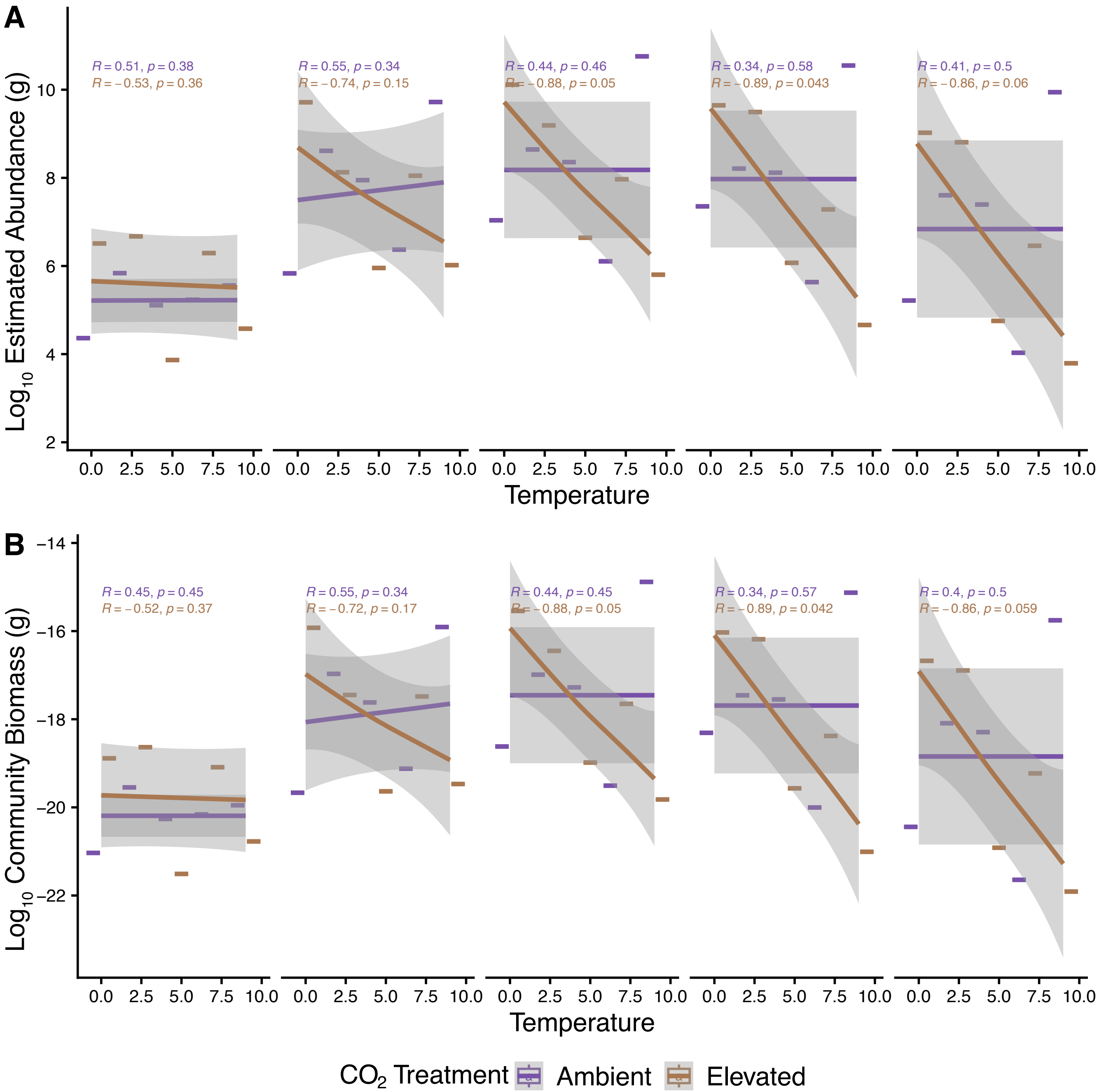


**Fig. S5.** Three-way interaction of demographic parameters. Significant empirical linear regressions for ambient $CO_2$ (purple lines) and elevated $CO_2$ (gold lines) across temperatures. Generalized additive model (GAM) regression bound by 95% confidence intervals (light gray). Estimated abundance (A) shows an increase across temperatures under ambient $CO_2$ with a slope modulated by size class and a decrease across temperatures under elevated $CO_2$ also accelerated by size class. (B) Community biomass shows the same pattern as estimated abundance (A), with a three-way interaction, as the main component of the overall biomass is abundance, due to the smaller relative changes in mean volume compared to mean changes in total number of individuals (Figs. 2, 3).

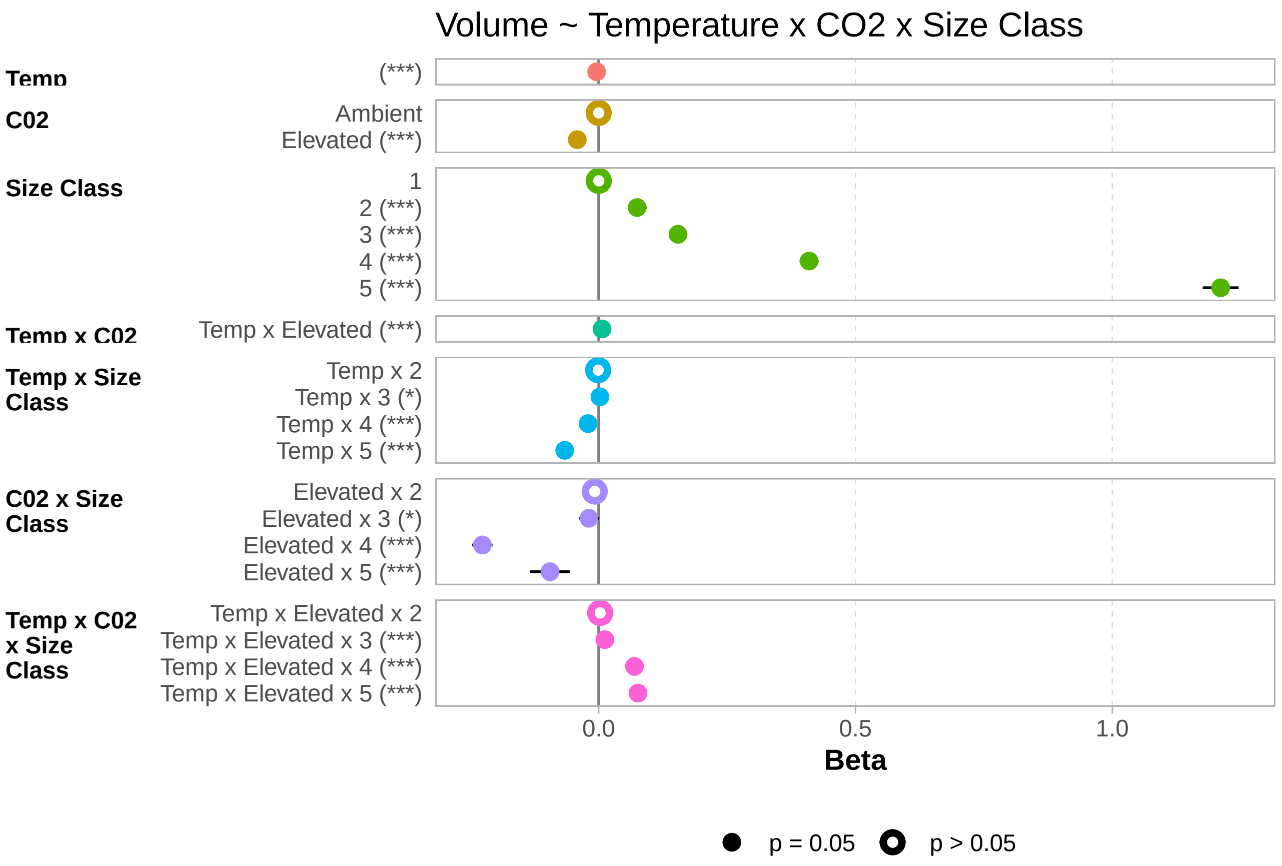


**Fig. S6.** Weighted regression slope $\beta$ coefficients for three-way interaction GLM model of volume. Model equation in title. Reference factor level stated. Significance levels: (*) $p \leq 0.05$; (**) $p \leq 0.01$; (***) $p \leq 0.001$.

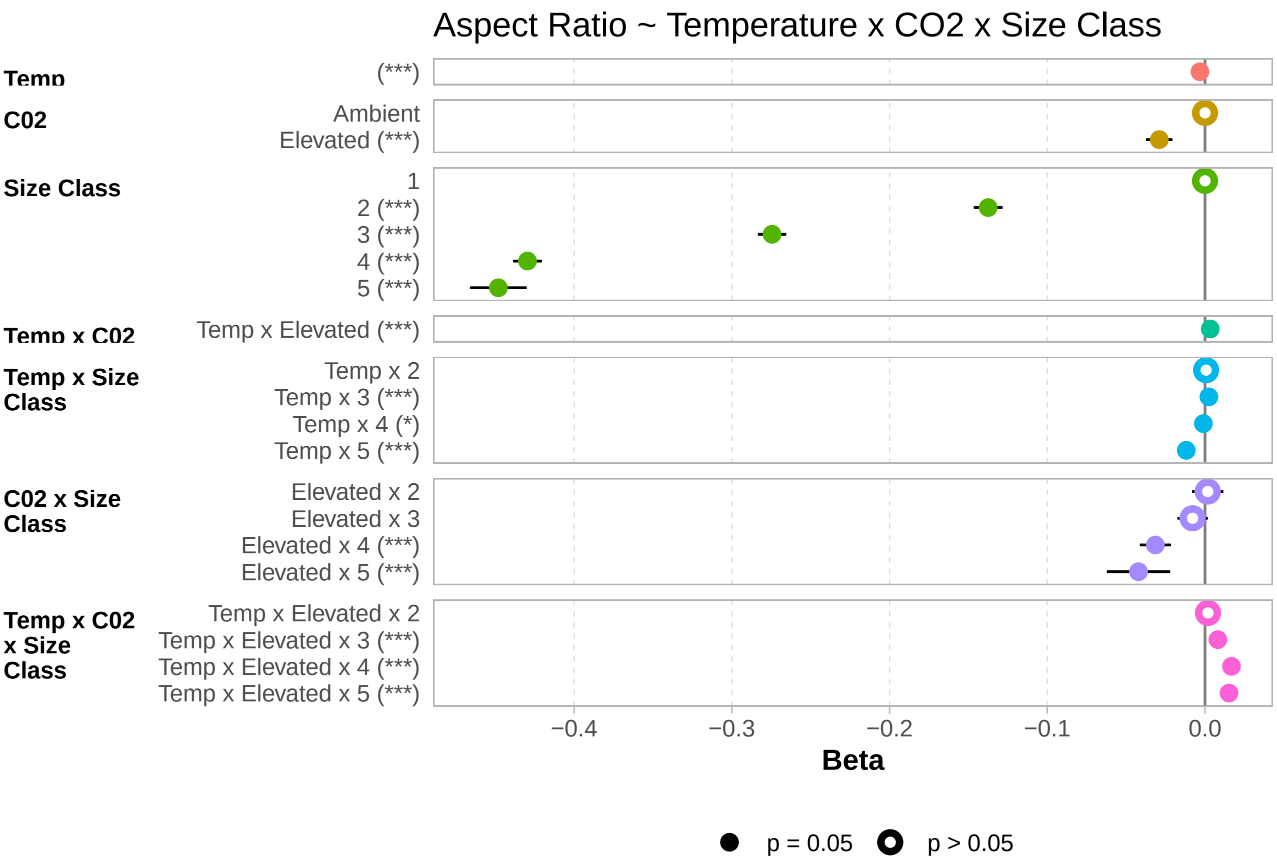


**Fig. S7.** Weighted regression slope $\beta$ coefficients for three-way interaction GLM model of aspect ratio. Model equation in title. Reference factor level stated. Significance levels: (*) $p \leq 0.05$; (**) $p \leq 0.01$; (***) $p \leq 0.001$.

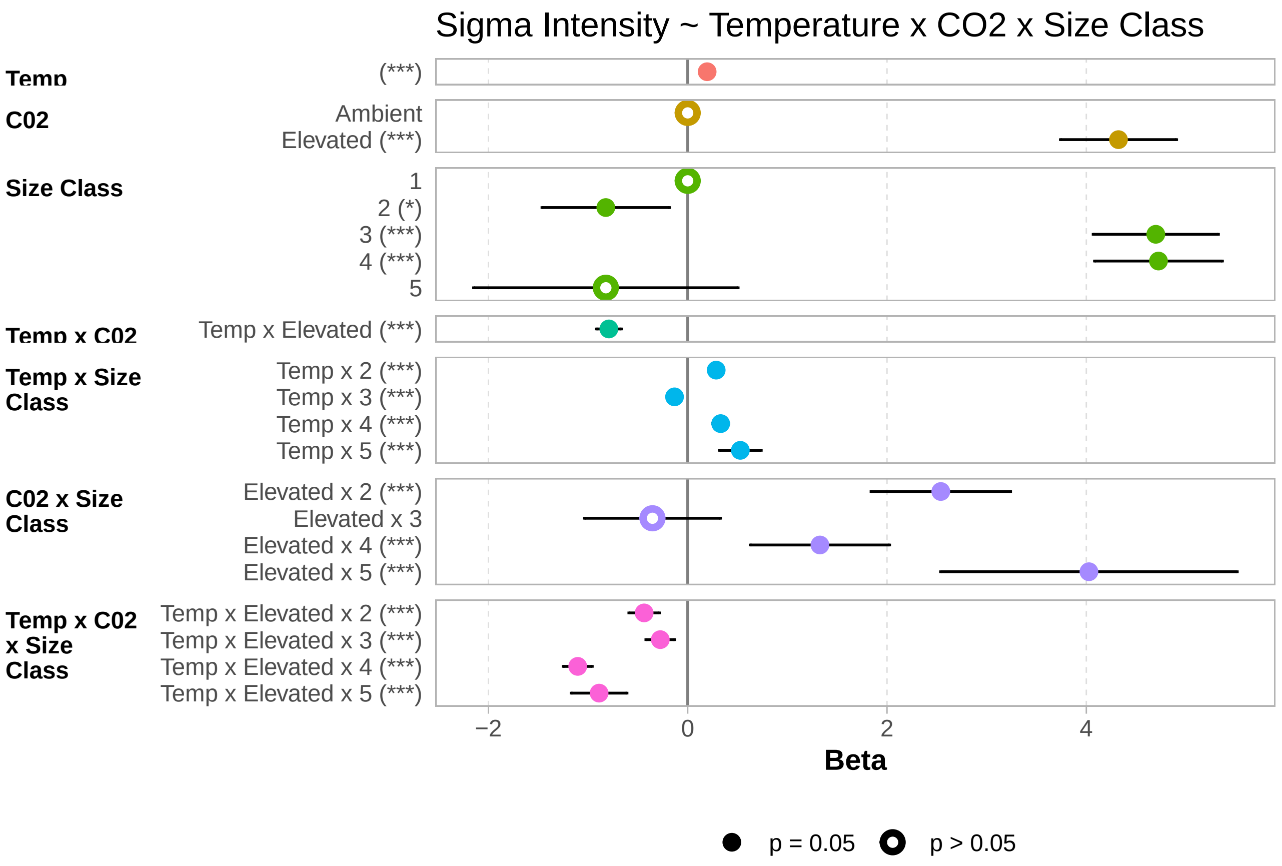


**Fig. S8.** Weighted regression slope $\beta$ coefficients for three-way interaction GLM model of sigma intensity. Model equation in title. Reference factor level stated. Significance levels: (*) $p \leq 0.05$; (**) $p \leq 0.01$; (***) $p \leq 0.001$.

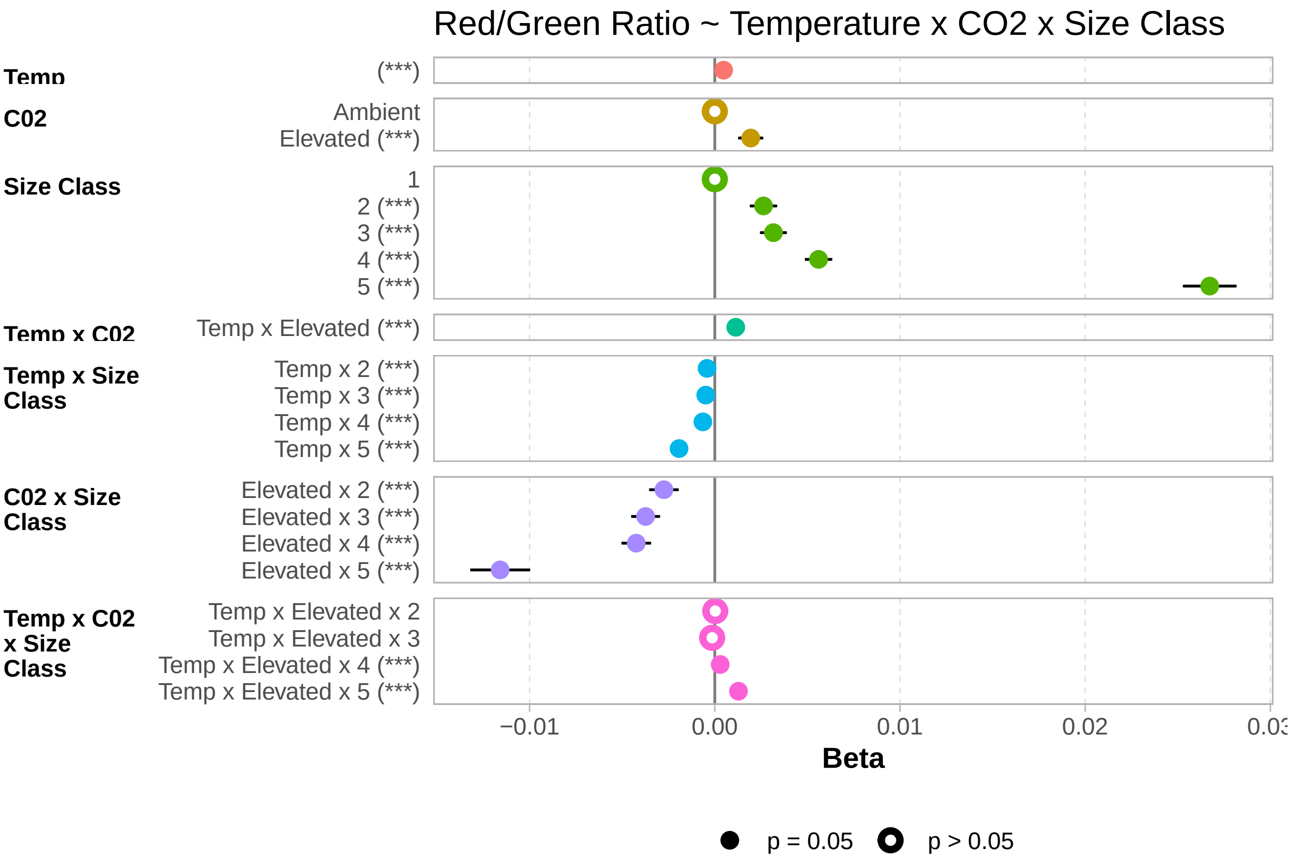


**Fig. S9.** Weighted regression slope $\beta$ coefficients for three-way interaction GLM model of red/green ratio. Model equation in title. Reference factor level stated. Significance levels: (*) $p \leq 0.05$; (**) $p \leq 0.01$; (***) $p \leq 0.001$.

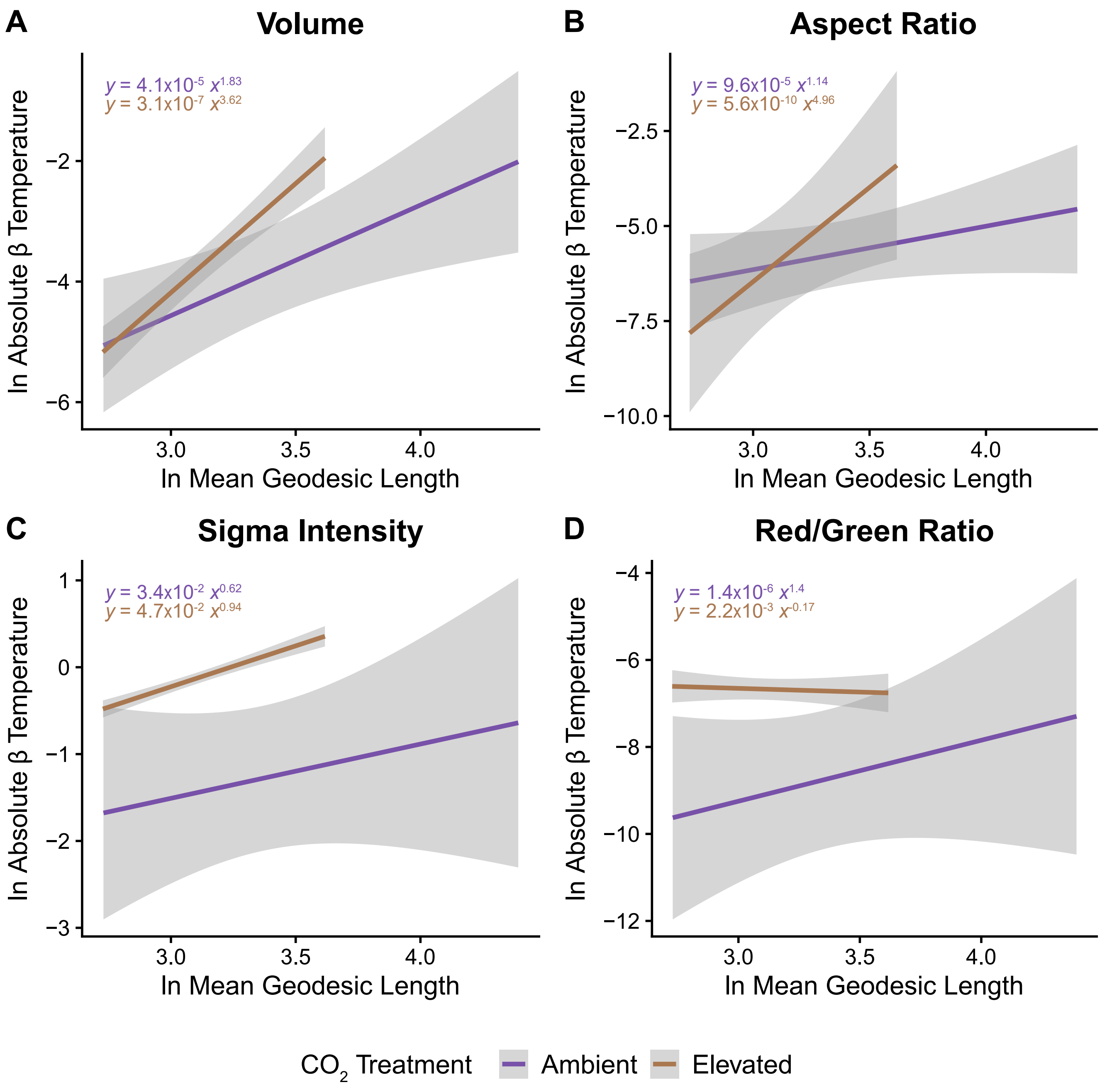


**Fig. S10.** Allometric relationship to determine size-dependent rates of change in functional traits by log-log plot: natural log of the absolute slope derived from GLMs for each size class and $CO_2$ treatment (Figs. 3; S6-S9) versus the natural log of the size class' mean geodesic length. Non-logged equation in corner. (A) Volume displays a strong super-linearity for ambient ($p = 0.06$) and significant super-linearity for elevated ($p = 0.01$). Support for size-dependency exists under elevated $CO_2$ for (C) sigma intensity ($p = 0.01$). All other relationships, though different from linear, were non-significant: (A) aspect ratio (0.215, 0.141), (B) sigma intensity under ambient $CO_2$, and (C) red/green ratio.

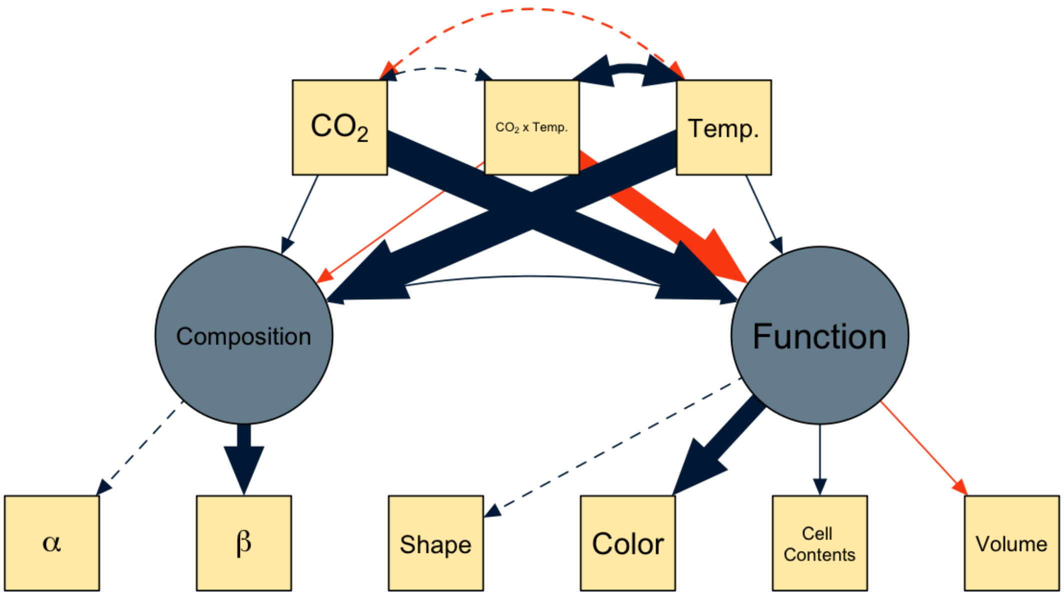


**Fig. S11.** Structural Equation Model (SEM) illustrating the effects of environmental variables on protist community structure and functional traits. Solid lines represent significant effects, with positive effects (blue) and negative effects (orange). The thickness of the lines corresponds to the magnitude of the standardized estimates. Dashed lines denote fixed latent variables without full parameter estimates (Table. S2). Nodes represent observed variables (yellow squares) and latent variables (blue circles). $\alpha$ represents total species richness, while $\beta$ represents the Shannon Diversity Index of treatments. Traits are as follows: Shape = Aspect Ratio, Color = Red to Green Ratio, Cell Contents = Sigma Intensity. Maximum Likelihood (ML) estimation converged after 328 iterations and the model provided a robust representation of our data ($\chi^2(20) = 121506.203$, $p < 0.001$). Despite the adequate convergence, the fit indices suggest that the model's ability to capture the complexity of ecological interactions could be improved (CFI = 0.832, TLI = 0.723, RMSEA = 0.197, SRMR = 0.074). Within the model's framework, a reversal of these effects was observed in the interaction between temperature and $CO_2$ (Estimate = -0.003, $p < 0.001$), which offset the influence of each factor alone (Table. S2).

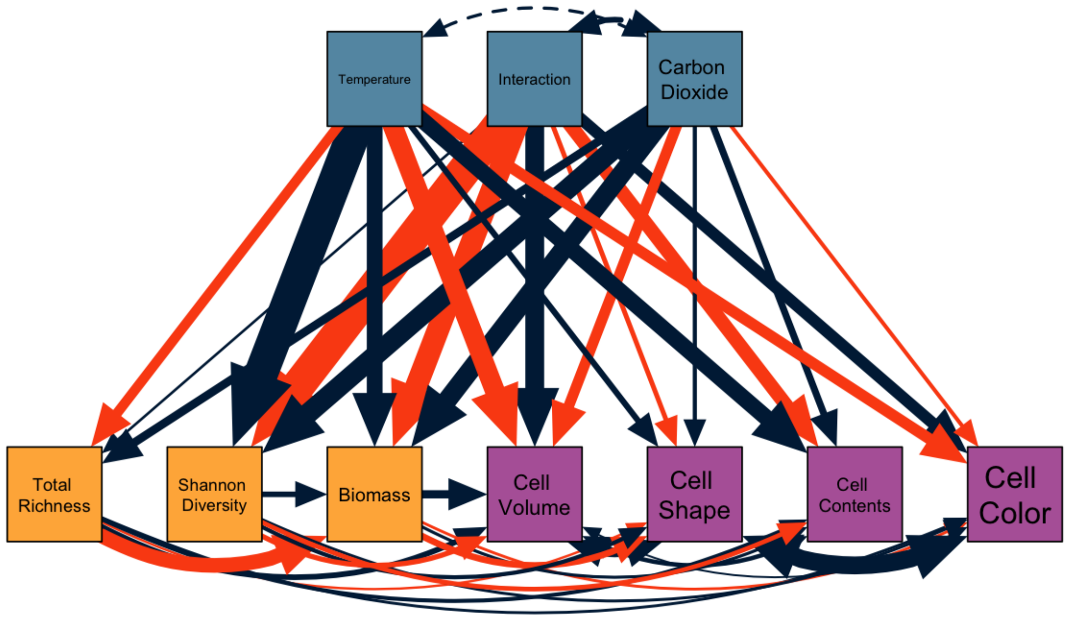


**Fig. S12.** Structural Equation Model (SEM) illustrating the effects of environmental variables (light blue) on protist community structure (light orange) and functional traits (purple). Solid lines represent significant effects, with positive effects (blue) and negative effects (red). The thickness of the lines corresponds to the magnitude of the standardized estimates. Dashed lines denote exogenous variables (Table. S3). Traits are as follows: Shape = Aspect Ratio, Color = Red to Green Ratio, Cell Contents = Sigma Intensity. Maximum Likelihood (ML) estimation converged after 337 iterations and the model provided a robust representation of our data ($\chi^2(20) = 58445.642$, $p < 0.001$). Along with adequate convergence, the fit indices suggest that the model captures well the complexity of ecological interactions (CFI = 0.958, TLI = -0.775, RMSEA = 0.61, SRMR = 0.025)

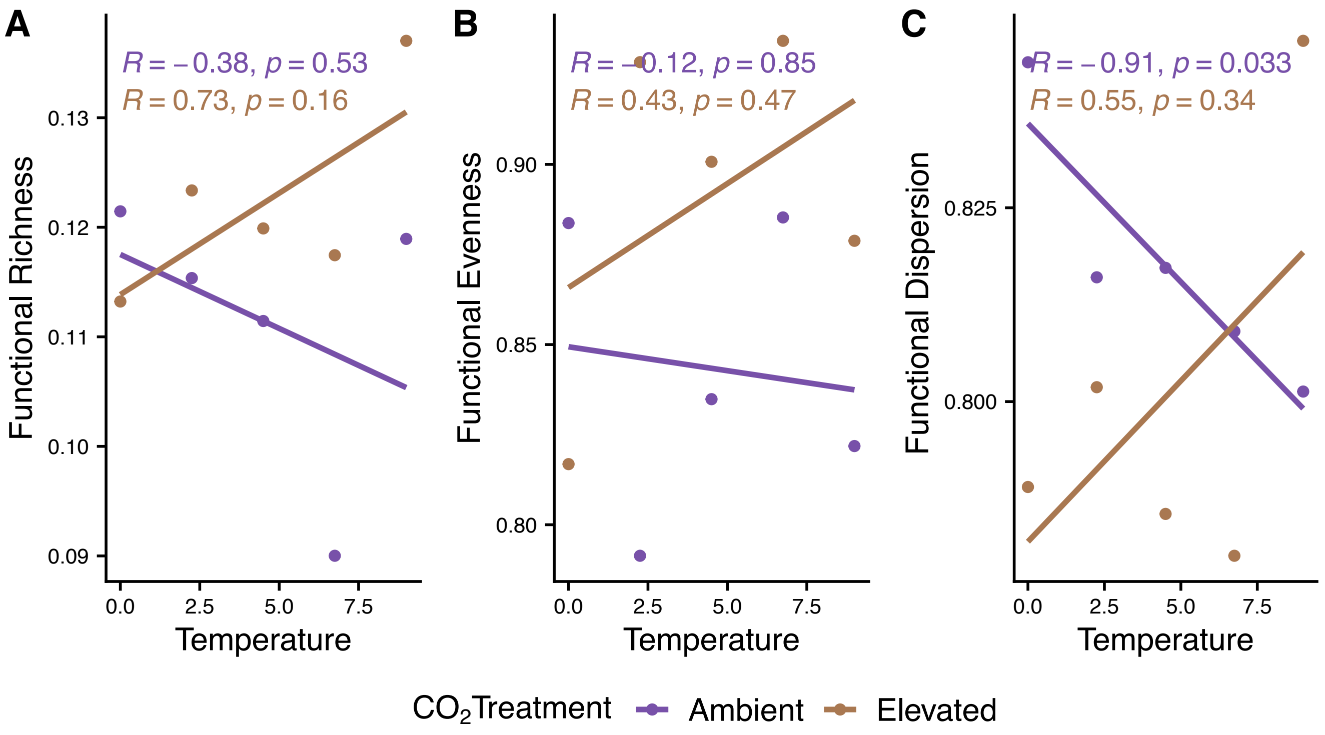


**Fig. S13.** Responses of Functional Traits to Temperature Under Different $CO_2$ Treatments. Panel A depicts the relationship between functional richness and temperature, showing a negative correlation for ambient $CO_2$ levels (purple line) and a positive correlation for elevated $CO_2$ levels (gold line). Panel B presents the correlation between functional evenness and temperature, indicating a non-significant relationship for both $CO_2$ treatments. Panel C illustrates a strong negative correlation between functional dispersion and temperature under ambient $CO_2$ levels, while the correlation under elevated $CO_2$ levels is not significant. The correlation coefficients (R) and their corresponding p-values are indicated for each trendline. Data points are represented as dots, with purple indicating ambient $CO_2$ conditions and gold indicating elevated $CO_2$ conditions.

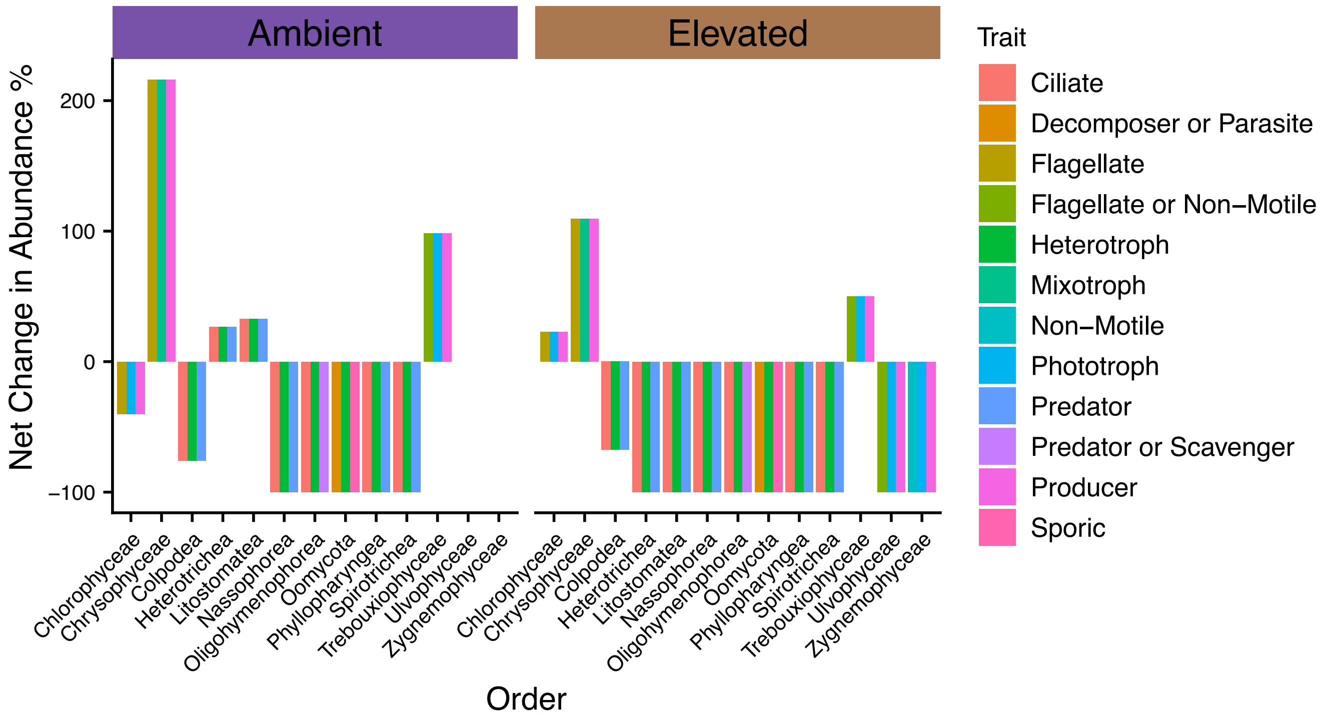


**Fig. S14.** Shifts in the relative abundance of Orders colored by functional trait group values from lowest to highest temperature treatment wrapped by carbon dioxide treatment (purple for ambient and gold for elevated). Overall, there is an increase in the relative abundance of phototrophic organisms and a decrease in the relative abundance of heterotrophs. Traits include resource acquisition strategy (heterotroph, mixotroph, autotroph), movement strategy (flagella, cillia, non-motile, sporic), and trophic level (producer, predator, decomposer, parasite, scavenger) taken from a literature search.

#### D. Tables.

**Table S1.** Transformation of traits to normalize distributions before analysis. Skewness calculated with R package *moments*. $\text{Log}_{10}$ transformation applied (denoted as **log** in table) with non-zero skewed data. Data containing zeroes transformed with square-root or geometric functions.

| Trait | Skewness | Distribution | Contains Zero | Transformation |
|---|---|---|---|---|
| Width | 3.56 | right-tailed | No | log |
| Volume (Derived) | 23.9 | right-tailed | No | log |
| Volume (ESD) | 50.39 | right-tailed | No | log |
| Volume (ABD) | 37.73 | right-tailed | No | log |
| Transparency | 1.08 | right-tailed | No | log |
| Symmetry | -1.26 | left-tailed | Yes | sqrt |
| Sum Intensity | 12.19 | right-tailed | No | log |
| Sigma Intensity | -0.34 | normal | No | N/A |
| Roughness | 3.83 | right-tailed | No | log |
| Ratio Red/Green | 2.99 | normal | No | N/A |
| Ratio Red/Blue | 1.50 | normal | No | N/A |
| Ratio Blue/Green | -0.82 | normal | No | N/A |
| Perimeter | 6.71 | right-tailed | No | log |
| Length | 6.41 | right-tailed | No | log |
| Intensity | -1.33 | normal | No | N/A |
| Geodesic Thickness | 2.84 | right-tailed | No | log |
| Geodesic Length | 6.55 | right-tailed | No | log |
| Geodesic Aspect Ratio | -0.01 | normal | No | N/A |
| Fiber Straightness | -0.61 | left-tailed | No | log |
| Fiber Curl | 3.03 | right-tailed | No | log |
| Feret Angle Minimum | 0.36 | circular | Yes | cotangent |
| Feret Angle Maximum | 0.39 | circular | Yes | cotangent |
| Elongation | 13.04 | right-tailed | No | log |
| Diameter (FD) | 5.94 | right-tailed | No | log |
| Diameter (ESD) | 5.99 | right-tailed | No | log |
| Diameter (ABD) | 5.74 | right-tailed | No | log |
| Convexity | -2.98 | left-tailed | No | log |
| Convex Perimeter | 5.99 | right-tailed | No | log |
| Compactness | 15.65 | right-tailed | No | log |
| Circularity Hu | -0.25 | normal | No | N/A |
| Circularity | -1.57 | left-tailed | No | log |
| Average Red | -1.43 | normal | No | N/A |
| Average Green | -1.43 | normal | No | N/A |
| Average Blue | -1.07 | normal | No | N/A |
| Aspect Ratio | 0.72 | bi-modal | No | N/A |
| Area Filled | 16.63 | right-tailed | No | log |
| Area (ABD) | 17.83 | right-tailed | No | log |

**Table S2.** Parameter estimates from the structural equation model analysis depicting the relationships between compositional diversity, functional diversity, and environmental variables. Each row represents a different pathway in the model, with the estimated coefficient, standard error, z-value, and p-value for each parameter. Significant pathways provide insights into how compositional and functional aspects of the community structure are influenced by temperature and carbon dioxide levels, as well as their interaction.

| | Left Hand Side | OP | Right Hand Side | Est. | S.E. | Z | p-value |
|---|---|---|---|---|---|---|---|
| 1 | CompositionalDiversity | =~ | Observed | 2.879 | 0 | | |
| 2 | CompositionalDiversity | =~ | Shannon | 0.134 | 0.0003 | 387.084 | 0 |
| 3 | FunctionalDiversity | =~ | Geodesic.Aspect.Ratio | 0.026 | 0 | | |
| 4 | FunctionalDiversity | =~ | Ratio.Red.Green | 0.109 | 0.002 | 52.723 | 0 |
| 5 | FunctionalDiversity | =~ | Sigma.Intensity | 0.199 | 0.004 | 44.308 | 0 |
| 6 | FunctionalDiversity | =~ | Volume | -0.073 | 0.001 | -53.210 | 0 |
| 7 | CompositionalDiversity | ~ | Temperature | 1.550 | 0.006 | 272.979 | 0 |
| 8 | CompositionalDiversity | ~ | Temperature:Carbon.Dioxide | -0.003 | 0.00001 | -301.992 | 0 |
| 9 | CompositionalDiversity | ~ | Carbon.Dioxide | 0.016 | 0.0001 | 299.607 | 0 |
| 10 | FunctionalDiversity | ~ | Temperature | 0.557 | 0.014 | 40.397 | 0 |
| 11 | FunctionalDiversity | ~ | Temperature:Carbon.Dioxide | -0.001 | 0.00002 | -39.718 | 0 |
| 12 | FunctionalDiversity | ~ | Carbon.Dioxide | 0.005 | 0.0001 | 41.783 | 0 |
| 13 | CompositionalDiversity | ~~ | FunctionalDiversity | 0.226 | 0.010 | 23.117 | 0 |
| 14 | Observed | ~~ | Observed | 67.997 | 0.256 | 265.938 | 0 |
| 15 | Shannon | ~~ | Shannon | -0.006 | 0.0002 | -31.534 | 0 |
| 16 | Geodesic.Aspect.Ratio | ~~ | Geodesic.Aspect.Ratio | 1.012 | 0.004 | 275.505 | 0 |
| 17 | Ratio.Red.Green | ~~ | Ratio.Red.Green | 1.202 | 0.006 | 186.427 | 0 |
| 18 | Sigma.Intensity | ~~ | Sigma.Intensity | 1.681 | 0.017 | 99.089 | 0 |
| 19 | Volume | ~~ | Volume | 1.091 | 0.005 | 240.735 | 0 |
| 20 | CompositionalDiversity | ~~ | CompositionalDiversity | 1.793 | 0.012 | 153.669 | 0 |
| 21 | FunctionalDiversity | ~~ | FunctionalDiversity | -17.436 | 0.599 | -29.125 | 0 |
| 22 | Temperature | ~~ | Temperature | 15.385 | 0 | | |
| 23 | Temperature | ~~ | Temperature:Carbon.Dioxide | 6,775.889 | 0 | | |
| 24 | Temperature | ~~ | Carbon.Dioxide | -691.407 | 0 | | |
| 25 | Temperature:Carbon.Dioxide | ~~ | Temperature:Carbon.Dioxide | 3,432,993.000 | 0 | | |
| 26 | Temperature:Carbon.Dioxide | ~~ | Carbon.Dioxide | -244,050.200 | 0 | | |
| 27 | Carbon.Dioxide | ~~ | Carbon.Dioxide | 49,125.440 | 0 | | |

**Table S3.** Parameter estimates from the structural equation model analysis depicting the relationships between compositional diversity, functional diversity, and environmental variables without latent variables. Each row represents a different pathway in the model, with the estimated coefficient, standard error, z-value, and p-value for each parameter. Significant pathways provide insights into how compositional and functional aspects of the community structure are influenced by temperature and carbon dioxide levels, as well as their interaction.

| | **Left Hand Side** | **OP** | **Right Hand Side** | **Est.** | **S.E.** | **Z** | **p-value** |
|---|---|---|---|---|---|---|---|
| 1 | Observed | ~ | Temperature | -0.311 | 1.075 | -0.289 | 0.772 |
| 2 | Observed | ~ | Temperature:Carbon.Dioxide | -0.003 | 0.002 | -1.830 | 0.067 |
| 3 | Observed | ~ | Carbon.Dioxide | 0.030 | 0.008 | 3.630 | 0.0003 |
| 4 | Shannon | ~ | Temperature | -0.048 | 0.055 | -0.866 | 0.386 |
| 5 | Shannon | ~ | Temperature:Carbon.Dioxide | -0.0001 | 0.0001 | -0.812 | 0.417 |
| 6 | Shannon | ~ | Carbon.Dioxide | 0.001 | 0.0004 | 2.136 | 0.033 |
| 7 | Geodesic.Aspect.Ratio | ~ | Temperature | -0.070 | 0.140 | -0.498 | 0.618 |
| 8 | Geodesic.Aspect.Ratio | ~ | Temperature:Carbon.Dioxide | 0.0001 | 0.0002 | 0.459 | 0.646 |
| 9 | Geodesic.Aspect.Ratio | ~ | Carbon.Dioxide | -0.0004 | 0.001 | -0.310 | 0.756 |
| 10 | Ratio.Red.Green | ~ | Temperature | -0.190 | 0.133 | -1.434 | 0.152 |
| 11 | Ratio.Red.Green | ~ | Temperature:Carbon.Dioxide | 0.0004 | 0.0002 | 1.874 | 0.061 |
| 12 | Ratio.Red.Green | ~ | Carbon.Dioxide | -0.001 | 0.001 | -1.002 | 0.316 |
| 13 | Sigma.Intensity | ~ | Temperature | 0.394 | 0.108 | 3.660 | 0.0003 |
| 14 | Sigma.Intensity | ~ | Temperature:Carbon.Dioxide | -0.001 | 0.0002 | -3.931 | 0.0001 |
| 15 | Sigma.Intensity | ~ | Carbon.Dioxide | 0.002 | 0.001 | 2.284 | 0.022 |
| 16 | Volume | ~ | Temperature | -0.106 | 0.139 | -0.765 | 0.444 |
| 17 | Volume | ~ | Temperature:Carbon.Dioxide | 0.0002 | 0.0002 | 0.770 | 0.441 |
| 18 | Volume | ~ | Carbon.Dioxide | -0.001 | 0.001 | -0.441 | 0.659 |
| 19 | Geodesic.Aspect.Ratio | ~ | Observed | -0.001 | 0.018 | -0.046 | 0.963 |
| 20 | Geodesic.Aspect.Ratio | ~ | Shannon | -0.080 | 0.355 | -0.224 | 0.823 |
| 21 | Ratio.Red.Green | ~ | Observed | -0.004 | 0.017 | -0.258 | 0.796 |
| 22 | Ratio.Red.Green | ~ | Shannon | 0.303 | 0.337 | 0.900 | 0.368 |
| 23 | Sigma.Intensity | ~ | Observed | 0.044 | 0.014 | 3.166 | 0.002 |
| 24 | Sigma.Intensity | ~ | Shannon | -0.411 | 0.273 | -1.504 | 0.132 |
| 25 | Volume | ~ | Observed | 0.006 | 0.018 | 0.340 | 0.734 |
| 26 | Volume | ~ | Shannon | -0.185 | 0.353 | -0.524 | 0.600 |
| 27 | Observed | ~~ | Observed | 58.500 | 11.700 | 5 | 0.00000 |
| 28 | Shannon | ~~ | Shannon | 0.154 | 0.031 | 5 | 0.00000 |
| 29 | Geodesic.Aspect.Ratio | ~~ | Geodesic.Aspect.Ratio | 0.973 | 0.195 | 5 | 0.00000 |
| 30 | Ratio.Red.Green | ~~ | Ratio.Red.Green | 0.878 | 0.176 | 5 | 0.00000 |
| 31 | Sigma.Intensity | ~~ | Sigma.Intensity | 0.577 | 0.115 | 5 | 0.00000 |
| 32 | Volume | ~~ | Volume | 0.964 | 0.193 | 5 | 0.00000 |
| 33 | Geodesic.Aspect.Ratio | ~~ | Ratio.Red.Green | -0.617 | 0.157 | -3.928 | 0.0001 |
| 34 | Geodesic.Aspect.Ratio | ~~ | Sigma.Intensity | -0.320 | 0.115 | -2.776 | 0.005 |
| 35 | Geodesic.Aspect.Ratio | ~~ | Volume | -0.773 | 0.175 | -4.410 | 0.00001 |
| 36 | Ratio.Red.Green | ~~ | Sigma.Intensity | -0.069 | 0.101 | -0.680 | 0.497 |
| 37 | Ratio.Red.Green | ~~ | Volume | 0.819 | 0.174 | 4.702 | 0.00000 |
| 38 | Sigma.Intensity | ~~ | Volume | 0.037 | 0.106 | 0.351 | 0.726 |
| 39 | Temperature | ~~ | Temperature | 10.125 | 0 | | |
| 40 | Temperature | ~~ | Temperature:Carbon.Dioxide | 6,834.375 | 0 | | |
| 41 | Temperature | ~~ | Carbon.Dioxide | 0 | 0 | | |
| 42 | Temperature:Carbon.Dioxide | ~~ | Temperature:Carbon.Dioxide | 6,150,938.000 | 0 | | |
| 43 | Temperature:Carbon.Dioxide | ~~ | Carbon.Dioxide | 227,812.500 | 0 | | |
| 44 | Carbon.Dioxide | ~~ | Carbon.Dioxide | 50,625 | 0 | | |

**Table S4.** Net change in relative abundance of Orders grouped by carbon dioxide treatment between +0 and +9 centigrade treatments, with relevant functional group information on resource acquisition, trophic level, and motility.

| Order | $CO_2$ | Net Change (%) | Nutritional Mode | Movement | Trophic Level |
|---|---|---|---|---|---|
| Chlorophyceae | Ambient | -40.208 | Phototroph | Flagellate | Producer |
| Chlorophyceae | Elevated | 22.769 | Phototroph | Flagellate | Producer |
| Chrysophyceae | Ambient | 216.054 | Mixotroph | Flagellate | Producer |
| Chrysophyceae | Elevated | 109.585 | Mixotroph | Flagellate | Producer |
| Colpodea | Ambient | -75.878 | Heterotroph | Ciliate | Predator |
| Colpodea | Elevated | -67.580 | Heterotroph | Ciliate | Predator |
| Heterotrichea | Ambient | 26.569 | Heterotroph | Ciliate | Predator |
| Heterotrichea | Elevated | -100 | Heterotroph | Ciliate | Predator |
| Litostomatea | Ambient | 32.855 | Heterotroph | Ciliate | Predator |
| Litostomatea | Elevated | -100 | Heterotroph | Ciliate | Predator |
| Nassophorea | Ambient | -100 | Heterotroph | Ciliate | Predator |
| Nassophorea | Elevated | -100 | Heterotroph | Ciliate | Predator |
| Oligohymenophorea | Ambient | -100 | Heterotroph | Ciliate | Predator or Scavenger |
| Oligohymenophorea | Elevated | -100 | Heterotroph | Ciliate | Predator or Scavenger |
| Oomycota | Ambient | -100 | Heterotroph | Sporic | Decomposer or Parasite |
| Oomycota | Elevated | -100 | Heterotroph | Sporic | Decomposer or Parasite |
| Phyllopharyngea | Ambient | -100 | Heterotroph | Ciliate | Predator |
| Phyllopharyngea | Elevated | -100 | Heterotroph | Ciliate | Predator |
| Spirotrichea | Ambient | -100 | Heterotroph | Ciliate | Predator |
| Spirotrichea | Elevated | -100 | Heterotroph | Ciliate | Predator |
| Trebouxiophyceae | Ambient | 98.215 | Phototroph | Flagellate or Non-Motile | Producer |
| Trebouxiophyceae | Elevated | 50.214 | Phototroph | Flagellate or Non-Motile | Producer |
| Ulvophyceae | Ambient | 0 | Phototroph | Flagellate or Non-Motile | Producer |
| Ulvophyceae | Elevated | -100 | Phototroph | Flagellate or Non-Motile | Producer |
| Zygnemophyceae | Ambient | 0 | Phototroph | Non-Motile | Producer |
| Zygnemophyceae | Elevated | -100 | Phototroph | Non-Motile | Producer |
